# Elucidation of the mechanism of byproduct DNA synthesis in whole-transcript amplification by poly-A tagging and development of its specific suppression

**DOI:** 10.1101/2025.08.19.670999

**Authors:** Yohei Sasagawa, Yoshimi Iwayama, Minoru Yano, Itoshi Nikaido

## Abstract

Poly-A tagging is a whole-transcript amplification method that uses terminal transferase to add poly-A to single-stranded cDNA in a non-template-dependent manner, and is used in highly sensitive single-cell RNA-seq. However, this method has long been reported to amplify byproduct DNA derived from reverse-transcription primers, particularly under conditions of low RNA abundance, leading to experimental complexity and reduced accuracy in quantifying gene expression. In this study, we elucidated the mechanism by which byproduct DNA is amplified at the molecular level and developed a new strategy of suppressing it named Suppression Tagging (SupTag). SupTag makes use of TdT, which has extremely low extension efficiency at the 3′ recessive end, to stabilize dA/dT base pairs and inhibit poly-A extension of oligo-dT primers, thereby preventing the formation of precursor DNA for byproduct DNA. We demonstrated that SupTag can specifically suppress byproduct DNA using chemicals such as tetramethylammonium chloride (TMAC), modified nucleotides (e.g., 2F-dATP), and Super T-modified reverse-transcription primers. SupTag improves usability by streamlining purification processes and prevents reductions in sequence read output and quantitative sensitivity caused by byproduct DNA. Additionally, SupTag demonstrated the ability to suppress byproduct DNA amplification without using suppression PCR and is compatible with cDNA amplification introducing different adapter sequences at both ends. This study presents a novel suppression method based on the mechanism of amplification of byproduct DNA in the poly-A tagging method, providing a practical technology enabling high-precision transcriptomic analysis from low-abundance RNA.

## Introduction

Transcriptomic analysis has made significant contributions to the estimation of cell functions, the elucidation of disease mechanisms, and the exploration of new drug targets by comprehensively capturing the mRNA expressed within cells [1]. In recent years, the development of single-cell RNA sequencing (RNA-seq) and spatial transcriptomics has made it possible to detect cell-type-specific features and spatial context dependences that were previously averaged out in bulk analyses [2]. In these applications, whole-transcript amplification (also known as cDNA amplification) is essential before preparing a DNA sequencing library in order to analyze extremely small amounts of mRNA derived from a single cell or a small localized region in a tissue section using a sequencer [3][4][5][6][7][8][9][10][11][12]. This whole-transcript amplification (WTA) involves converting mRNA into amplifiable cDNA and amplifying the cDNA [13]. In these WTA processes that target mRNA, non-target sequences, also known as artifact or byproduct DNA, are synthesized through different mechanisms for each WTA process [5][8][14][15][16][17][18][19][20]. It has been reported that these phenomena can reduce the utilization rate of sequence reads and the capacity to accurately and sensitively quantify gene expression. Resolving the nucleic acid synthesis of these byproducts is generally important for obtaining high-quality gene expression profiles [21].

The work described in this paper is aimed at developing a method of suppressing byproduct DNA in the poly-A tagging method of WTA, which uses terminal transferase [22][23]. This approach, established in the 1980s, was used in the first reported single-cell RNA-seq experiment [24]. Terminal transferase, which is used in poly-A tagging, efficiently adds poly-A to nearly all single-stranded DNA molecules [25]. We present a schematic diagram showing the conventional mechanism of WTA using the poly-A tagging method described above (Fig. 1A). During reverse transcription, primers containing a 5′ adapter and 3′ oligo-dT sequence are used to generate single-stranded cDNA from poly-A RNA. Barcoded primers enable high-throughput processing. After purification, terminal transferase adds a poly-A tail to the 3′ end of the cDNA, a process referred to as poly-A tailing, to which a tagging primer anneals for second-strand synthesis. The resulting cDNA is amplified by PCR and used for sequencing. Amplified cDNA ranges from 450 to 5,000 bp in size (Fig. S1) [8][26]. Single-cell RNA-seq methods using poly-A tagging, such as Quartz-Seq and Quartz-Seq2, have achieved high sensitivity [8][26]. In a 2020 benchmark study comparing single-cell RNA-seq methods, Quartz-Seq2—the only method using poly-A tagging—detected 1.5–5 times more genes than the others [27].

**Figure 1.**
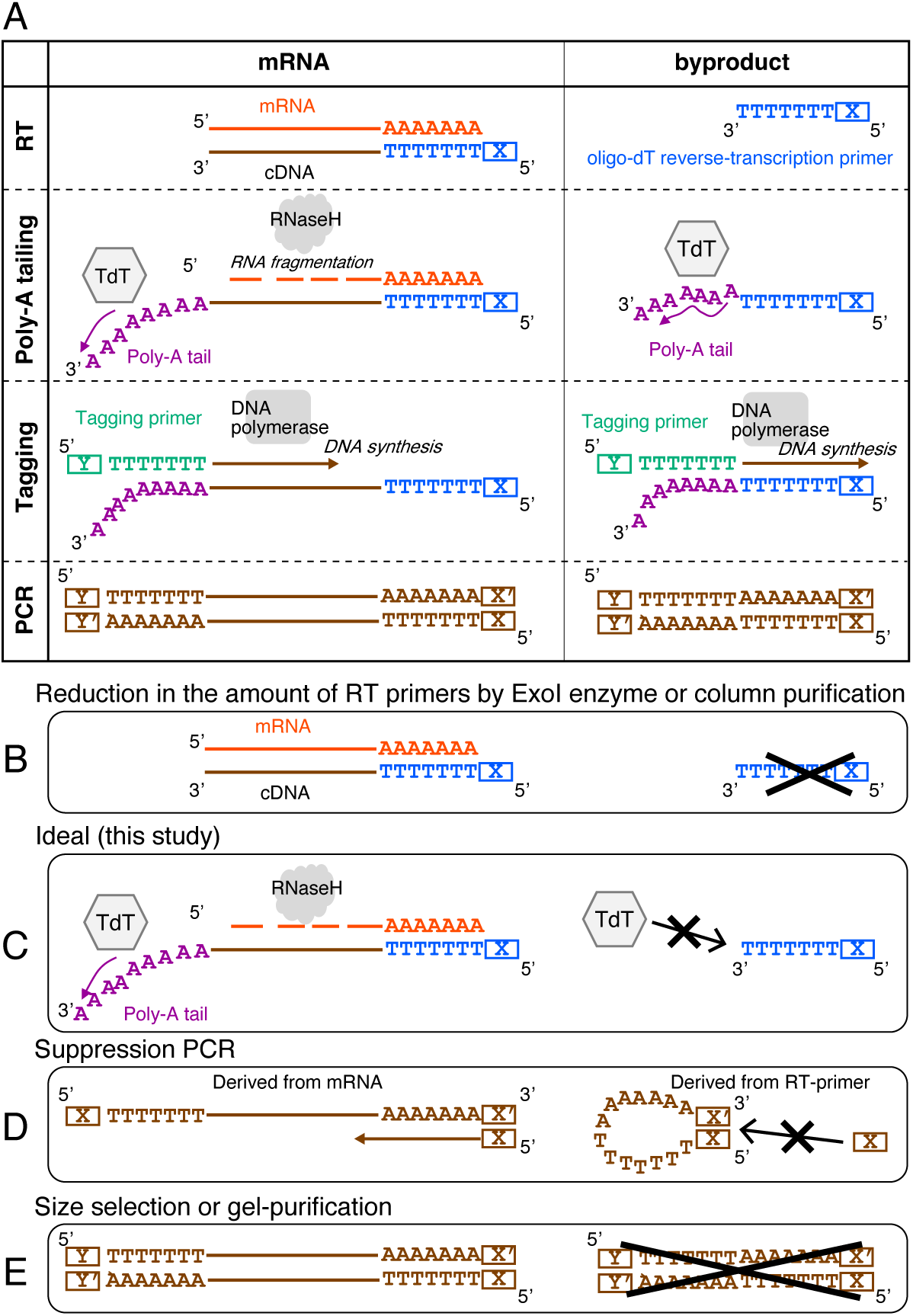
**A schematic representation of whole-transcript amplification using the poly-A tagging method**. The upper left panel shows a schematic representation of the molecular mechanism by which cDNA is amplified from mRNA during the process from reverse transcription to PCR amplification. The upper right panel shows the mechanism by which reverse-transcription primers are amplified as byproduct DNA by poly-A tagging. Panels A, C, and D represent previously reported strategies to reduce byproduct DNA derived from reverse-transcription primers. Panel B represents the strategy targeted in this study, aiming to specifically add poly-A to cDNA derived from mRNA while avoiding its addition to reverse-transcription primers.

However, the poly-A tagging method of WTA has the disadvantage of amplifying not only cDNA but also byproduct DNA derived from reverse-transcription primers, for which no effective solution has yet been proposed. Amplified byproduct DNA of poly-A tagging does not contain mRNA-derived sequences, and it has been reported that the presence and amplification of this byproduct DNA reduce the proportion of cDNA-derived sequences in the DNA sequencing library, thereby reducing the reproducibility, accuracy, and detection sensitivity of gene expression in transcriptomic analysis [19][20]. There is thus a need to remove byproduct DNA or suppress its amplification in order to minimize the adverse effects on transcriptomic analysis [8][20][24][26][28][29]. The impact of byproduct DNA from poly-A tagging is more pronounced when extremely small amounts of RNA are used as starting material [19][20]. Approximately 10¹¹ copies of reverse-transcription primers are used for a single cell, far exceeding the ∼10⁵ copies of mRNA present in a cell [8]. As a result, lower mRNA input increases the risk of amplifying byproduct DNA.

Next, we show the characteristics of byproduct DNA and the limitations of reported methods for dealing with it. Byproduct DNA from poly-A tagging typically ranges from 100 to 500 bp in size (Fig. S1C) [8]. While some of it can be removed via gel or bead purification, its complete elimination is difficult (Fig. 1E). In particular, the removal of large amounts of byproduct DNA makes library DNA preparation difficult due to the extended experimental times caused by the need for repeated purification and reduced cDNA yield [8][20][24][28][29]. Extending the reaction time of poly-A tailing increases the size of byproduct DNA, causing it to overlap with the cDNA size distribution and making its removal more difficult (Fig. S1). One approach for overcoming this problem is suppression PCR, which suppresses byproduct DNA amplification by adding identical adapters to both DNA ends, promoting pan-like structures after denaturation that block primer binding. This effect is stronger in short DNA fragments, selectively inhibiting short byproducts. However, this approach places constraints on the range of sequence library formats that can be used (Fig. 1D) [26]. Given these obstacles and the lack of an effective solution for specifically preventing the generation of precursors of amplifiable byproduct DNA in poly-A tagging, we developed a new method to address this issue (Fig. 1C).

In this study, we performed a detailed analysis of the mechanism of amplification of byproduct DNA in poly-A tagging and developed a new strategy called Suppression Tagging (SupTag) to suppress the synthesis of byproduct DNA based on the obtained findings. We demonstrated that SupTag suppresses the addition of poly-A to reverse-transcription primers under conditions that stabilize dA/dT base pairs. This effect is the result of specific inhibition of the amplification of byproduct DNA. Furthermore, SupTag enhances the stability, reproducibility, and reliability of WTA through the poly-A tagging method without compromising the amplification efficiency or sensitivity of cDNA. This is expected to contribute to improving the quality of transcriptomic analysis from low-abundance RNA.

## Results

### Detailed analysis of the effect of poly-A tailing on changes in the DNA size of reverse-transcription primers

To develop a novel method for specifically suppressing byproduct DNA generated by poly-A tagging, we first evaluated the characteristics of reverse-transcription primers that had been poly-A-tailed using terminal transferase. Poly-A-tailed reverse-transcription primers are molecules that serve as precursors for byproduct DNA. We performed poly-A tailing reactions or poly-T tailing reactions for 7 min on a 24-base oligonucleotide reverse-transcription primer that was conjugated with the fluorescent dye fluorescein (FAM) at the 5′ end. Subsequently, we analyzed the size distribution at single-base resolution. In the absence of dNTP, peaks were exclusively observed at the position corresponding to the 24 bases (Fig. 2A and Fig. S2). Under conditions with poly-T tailing, reverse-transcription primers exhibited a single-peak distribution exceeding 129 bases, indicating that almost all of the reverse-transcription primers contained at least 100 thymine bases and this trend was consistently observed in all four technical replicates (n=4). Even under conditions where poly-A tailing was performed, no signal was observed at the original region of 24 bases. In addition, the reverse-transcription primers showed a bimodal size distribution, with a sharp peak and a very broad peak observed around 48 bases (Fig. 2 and Fig. S2), as consistently observed in all four technical replicates. The position at 48 bases corresponds to the addition of 24 poly-A tails, which was considered to divide the signal into two groups: a sharp peak and a broad peak. Additionally, it was observed that the maximum length due to poly-A tailing was longer than that due to poly-T tailing. These results suggest that the oligo-dT primer contains a subgroup that is not easily targeted by terminal transferase for poly-A tailing.

**Figure 2.**
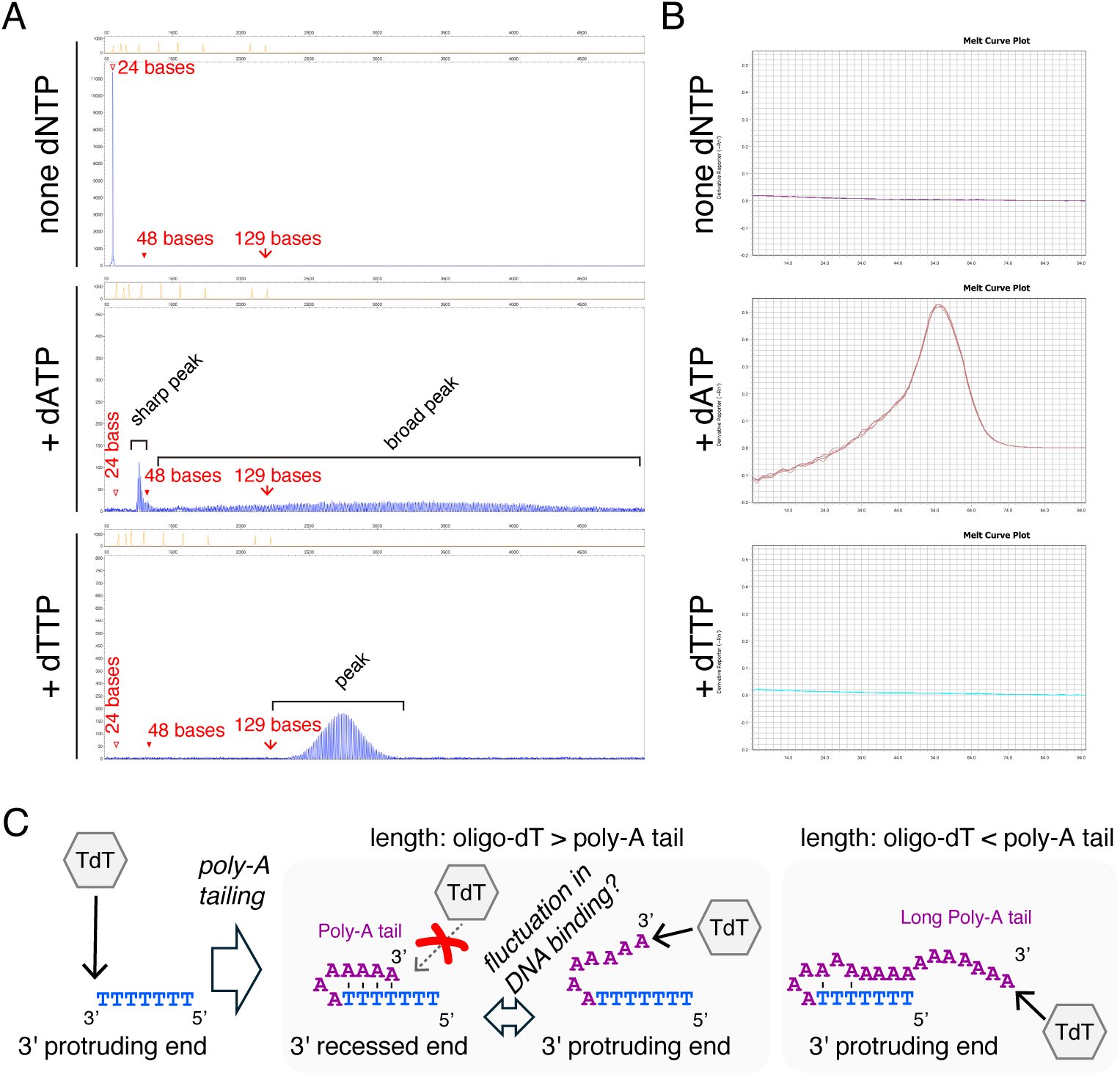
Characterization of reverse-transcription primers with poly-A tailing. a) A terminal transferase-mediated non-template-dependent base addition reaction was performed using dATP or dTTP on a 24-base oligo-dT primer modified with FAM at the 5′ end. Four technical replicates (n=4) were prepared and the typical pattern is shown. All size distribution profiles are shown in Fig. S2. The x-axis represents DNA size based on electrophoresis mobility. The y-axis shows the detection of DNA quantity for the LIZ120 size standard in orange and the detection of FAM fluorescence wavelength in blue. The positions of 24 bases, 48 bases, and 129 bases are indicated. b) After performing a TdT reaction on an oligo-dT primer without FAM labeling at the 5′ end, melting curve analysis was performed in five technical replicates (n=5). The x-axis indicates the melting curve temperature, and the y-axis indicates the fluorescence intensity. c) Hypothesis of the 3′-end structure when poly-A tailing was performed on reverse-transcription primers containing oligo-dT sequences. When the length of the oligo-dT primer is shorter than the length of the poly-A tail, self-annealing is expected to occur, forming a stem-loop structure. In this case, a 3′ recessed end structure is formed. Because dA/dT base pairs have a low annealing temperature, when the stem-loop structure breaks, it forms a 3′ protruding end. It is assumed that dA/dT base pairs transition between these two states. When the oligo-dT primer is longer than the poly-A tail, it is predicted to form a 3′ protruding end.

Subsequently, we proceeded to measure alterations in the DNA size distribution of reverse-transcription primers over time during the poly-A tailing reaction. We then analyzed the formation of the bimodal population resulting from the poly-A tailing reaction. Given that our previous observations had only been under 7-min reaction conditions, we measured changes in the size of the oligo-dT primer under shorter reaction times of 3, 4, and 5 min (Fig. S3). In the 3-min poly-A tailing reaction, poly-A was added to almost all oligo-dT primers, but the DNA size distribution showed a single peak in the range of 39–48 bases. Meanwhile, under 4- and 5-min reactions, DNA populations longer than 48 bases appeared, and under the 7-min reaction, these populations became even more elongated. Under the 5- and 7-min reactions, DNA populations with sharp peaks of 48 bases or less did not disappear, and a bimodal distribution was observed (Fig. S3). Interestingly, the shortest signal change in the sharp peak under the 3-min and 7-min reactions differed by only one base, from 39 to 40 bases. These results suggest that poly-A tailing causes almost all of the oligo-dT primers to be poly-A tailed, forming a single peak at 39–48 nt, and that, over time, some of this DNA undergoes elongation, resulting in a double peak. Additional experiments suggested that, following poly-A tailing of the oligo-dT primer, some of the single-peak DNA population with 48 bases or fewer has difficulty incorporating adenine bases.

### Melting curve analysis of poly-A-tailed oligo-dT primers

Next, we investigated the effects of differences in the sequence of the template DNA on non-template-dependent base addition caused by poly-A tailing/poly-T tailing. In particular, we investigated changes in the size corresponding to the peak with the highest-intensity signal (Fig. S4). Our results showed that, when using an oligo-dA primer as the template DNA, the position of the peak with the highest intensity for the poly-T tailing corresponded to a size 213 bases shorter than that for the poly-A tailing (poly-A tailing: 268.5±1.2 bases [n=4]; poly-T tailing: 55.5±1.0 bases [n=4]). When an oligo-dT primer was used as the template DNA, the position of the highest-intensity peak for the poly-A tailing was approximately 70 bases shorter than that for the poly-T tailing (poly-A tailing: 40.0±0.0 bases [n=4]; poly-T tailing: 110.2±0.9 bases [n=4]). Both the poly-A-containing oligo-dT primer and the poly-T-containing oligo-dA primer have a structure in which the poly-A and poly-T sequences are adjacent to each other. Under these conditions, fewer bases could be added by terminal transferase.

As described above, we decided to investigate whether the primer forms a double-stranded DNA structure under conditions where TdT is prevented from adding bases. We investigated whether the primers formed double-stranded DNA structures after the TdT reaction by performing a melting curve analysis with a double-strand-specific fluorescent dye (Fig. 2B). When the TdT reaction was performed without adding dNTPs, the melting curve of the oligo-dT primer was flat, suggesting that it did not form a double-stranded DNA structure. The same result was observed when dTTP was added. Meanwhile, TdT reaction products with dATP addition consistently showed a single peak, confirming the presence of a double-stranded DNA structure in five out of five cases. These results imply that the poly-A-tailed, single-stranded oligo-dT primer can form an intramolecular stem-loop structure (also known as a hairpin structure), locally forming complementary double strands composed of dA and dT.

### Hypothesis on the detailed mechanism of amplification of byproduct DNA by poly-A tagging

Based on the above results, we hypothesized that the following reactions occur during the poly-A tagging of WTA (Fig. 2C). When poly-A tailing is performed on an oligo-dT primer, short poly-A sequences are added to the 3′ terminal of the poly-T sequence. In this particular instance, a stem-loop structure (hairpin structure) can form between the poly-T sequence and the poly-A sequence. As previously reported, when single-stranded DNA forms a stem-loop structure, the loop length must be at least 3 bases, and 4 to 8 bases are required for stability [30]. Furthermore, when the length of added poly-A is shorter than that of oligo-dT, the terminal structure is expected to become a 3′ recessive end. Non-template-dependent base addition by terminal transferase has been reported to occur very slowy at the 3′ recessive end [31]. Meanwhile, the stability of base-pairing between dA and dT bases is lower than that between dG and dC bases. Consequently, it is anticipated that a transition between the stem-loop structure and single-stranded DNA states will occur. The dissociation between dA and dT bases results in protrusion of the 3′ end of single-stranded DNA, thereby facilitating poly-A tailing through the TdT reaction. When the poly-A tail becomes sufficiently long compared with the length of the oligo-dT primer, this state of protrusion of the 3′ end is more easily maintained, and TdT further extends the poly-A tail (Fig. 2C). In principle, DNA with a long poly-A tail is expected to bind more easily to the tagging primer and thus be amplified as byproduct DNA.

Based on the above hypothesis, we conceived the idea of performing poly-A addition by terminal transferase under conditions where base-pairing between dA and dT bases is stabilized as a method to specifically suppress the amplification of byproduct DNA caused by poly-A tagging derived from reverse-transcription primers. Assuming that the stability of the base-pairing between dA and dT bases can be improved, it is expected that the oligo-dT primer will maintain its stem-loop structure and 3′ recessive end, thereby facilitating the inhibition of poly-A tailing by terminal transferase. If the poly-A tail remains short, the tagging primer will have reduced affinity for binding to the same site. This will reduce the number of template molecules that are amplified as byproducts during PCR. Mammalian mRNA has a cap structure at the 5′ end and a poly(A) tail at the 3′ end [32]. However, to the best of our knowledge, no reports indicating that polyadenylation occurs at the 5′ end of mRNA have been published. Therefore, it is unlikely that features such as an oligo-dT sequence are present at the 3′ end of full-length cDNA corresponding to the 5′ end of mRNA. Additionally, the 3′ end of single-stranded cDNA after reverse transcription forms an RNA/DNA hybrid structure, which differs from the 3′ end of the reverse-transcription primer’s oligo-dT sequence, forming a single-stranded DNA (ssDNA) structure. As such, the 3′ ends of cDNA and reverse-transcription primers differ in both nucleotide sequence and the presence or absence of complementary base-pairing. Therefore, we considered that the proposed method described above could specifically suppress the amplification of byproducts (Fig. 1C).

### Effect of cDNA and byproduct DNA amplification on factors stabilizing dA/dT base pairs

Based on the method proposed above, we prepared candidate factors reported to stabilize base pairs between dA and dT and investigated whether byproduct-specific DNA amplification inhibition was possible. To do this, it was necessary to simultaneously measure the amounts of amplified cDNA and byproducts. We performed WTA using the poly-A tagging method with total RNA and quantified the size distribution and DNA quantity, with reference to previous methods [8]. In particular, the reverse-transcription reaction was carried out in PCR tubes instead of a 384-well plate, increasing the number of experiments that could be conducted in parallel (see Experimental Procedure). This enabled verification of the effects of poly-A tailing under various conditions on WTA. The analysis of byproduct DNA revealed a size distribution ranging from 100 base pairs (bp) to over 500 bp, while that of amplified cDNA was 450 bp or more (Fig. S1). In certain instances, the range of 450–500 bp may be observed to overlap between byproducts and cDNA. To avoid quantification in this overlapping range, the range of 100–450 bp was designated as byproduct DNA, while the range of 550–9000 bp was designated as cDNA. The amounts of each type of DNA were then quantified. First, the amount of amplified cDNA was quantified and the extent to which it changed upon applying each condition was evaluated. Next, the ratio of the amount of byproduct DNA to the amount of cDNA was used to evaluate the extent of change across different conditions as a Z-score (the deviation from the mean divided by the standard deviation). A small ratio indicates that the byproduct is specifically amplified and suppressed compared with cDNA; as such, a lower Z-score reflects a better result. Ideally, conditions that do not affect the quantity of amplified cDNA and result in a decrease in the Z-score are desirable.

First, we planned an experiment to add modified nucleotides to the poly-A tail (see Fig. S5A for details on the modified nucleotides). In the first experiment, WTA (technical replicates: n=3) was performed by replacing 25% of dATP with five modified nucleotides, and cDNA amplification was observed under all conditions (Fig. S5C). The five modified nucleotides used were as follows: dZTP (2-amino-2′-deoxyadenosine 5′-triphosphate), NH2dATP (2′-amino-2′-deoxyadenosine 5′-triphosphate), AzdATP (2′-azido-2′-deoxyadenosine 5′-triphosphate), OmdATP (2′-O-methyladenosine 5′-triphosphate), and 2F-dATP (2′-fluoro-2′-deoxyadenosine triphosphate) (Fig. S5A). 2F-dATP and OmdATP have been reported as factors that can stabilize the dA/dT base pairs [33][34]. Under conditions with 100% dATP, byproduct DNA was observed and the Z-score was consistently high (Fig. S5B). In contrast, under conditions with the addition of 2F-dATP, the Z-score was the lowest (Fig. S5B). Under OmdATP conditions, a tendency toward lower Z-scores was also observed (Fig. S5B). The amount of cDNA under 2F-dATP conditions was not significantly different from the average amount of cDNA under 100% dATP conditions (n=3, 82.7±23.6%). The amount of cDNA under OmdATP conditions was comparable to that under 100% dATP conditions (n=3, 95.7±3.1%). To confirm the reproducibility of these results, a second experiment was performed under the same conditions (Fig. 3 and Fig. S6). The results repeatedly revealed decreases in Z-score under the 2F-dATP and OmdATP conditions (Fig. 3B). As an additional condition, an experiment using 10% 2F-dATP was carried out, and a decrease in Z-score was observed, but this decrease was less pronounced than that observed under the 25% conditions. The cDNA levels under the 25% 2F-dATP and 10% 2F-dATP conditions were comparable to the average cDNA levels under the 100% dATP conditions (n=3: 95.6±4.3%, n=3: 99.9±2.5%). The amount of cDNA under the OmdATP conditions was comparable to that under the 100% dATP conditions (n=3, 94.9±4.6%). Thus, the addition of modified nucleotides such as 2F-dATP and OmdATP to the poly-A tail suppressed amplification specifically for byproduct DNA.

**Figure 3.**
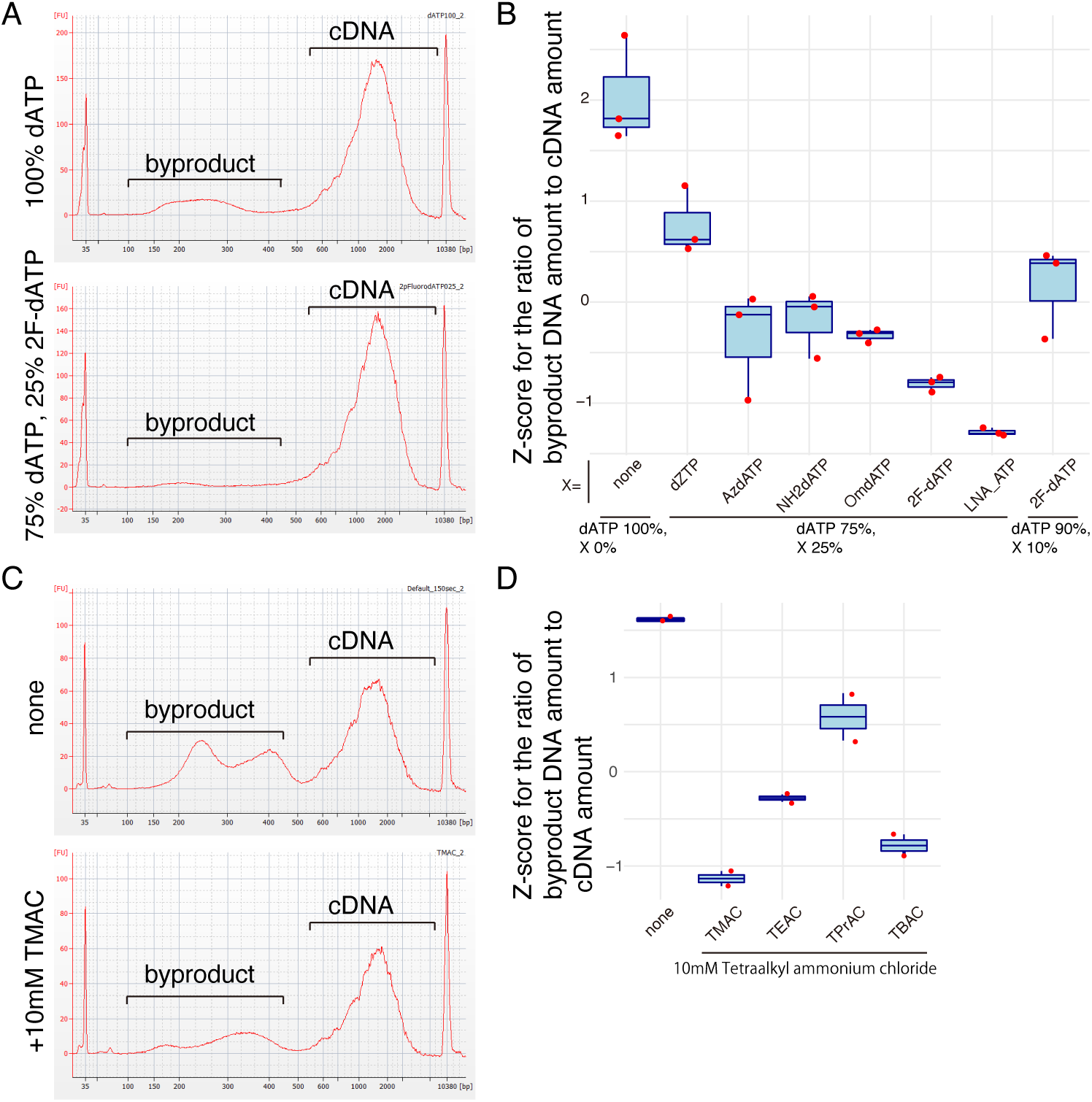
Effect of poly-A tagging on whole-transcript amplification of candidate factors stabilizing dA/dT base pairs. Byproduct DNA is classified as that of 100–450 bp, and amplified cDNA is classified as that of 550–9000 bp. The Z-score of the ratio of byproduct DNA to amplified cDNA was calculated and is shown. The reaction time for poly-A tailing was 75 s in (a) and (b), and 150 s in (c) and (d). a) shows the typical electrophoresis pattern of amplified cDNA in 2F-dATP. All patterns from technical replicates (n=3) are shown in Fig. S6. In b), the modified bases used and their mixing ratios are shown. The y-axis indicates the Z-score. In c), the typical electrophoresis pattern of amplified cDNA with 10 mM TMAC added is shown. All patterns, including those with other TAA compounds, are shown in Fig. S7. Fig. S7D shows the type of TAA compound used on the x-axis and the Z-score on the y-axis.

Next, we planned an experiment to add tetraalkylammonium chloride (TAA) to the poly-A tail (for details on this ammonium salt, see Fig. S7). We added 10 mM TAA to the poly-A tailing reaction and performed WTA (Fig. 3 and Fig. S7). We prepared five types of TAA: tetramethylammonium chloride (TMAC), tetraethylammonium chloride (TEAC), tetrapropylammonium chloride (TPrAC), tetrabutylammonium chloride (TBAC), and tetrapentylammonium chloride (TPeAC). It has been reported that TMAC and TEAC bind to dA/dT base pairs, thereby enhancing their stability [35]. Concentrations of TMAC below 100 mM have been reported to improve the specificity and amplification efficiency of PCR [36]. We amplified cDNA under these conditions and measured the amount of cDNA (see Fig. S7). The amount of cDNA under the three conditions (with TMAC, TEAC, and TPrAC) was comparable to the average amount of cDNA under the conditions without TAA addition (n=2: 97.2±4.9%, n=2: 102.9±12.5%, n=2: 107.1±11.7%). The amount of cDNA under TBAC conditions was slightly reduced compared with the average amount of cDNA under conditions without TAA addition (n=2: 76.2±1.69%). Under TPeAC conditions, no cDNA was amplified (Fig. S7). Meanwhile, under conditions without TAA addition, byproduct DNA was observed and the Z-score was generally high (Fig. 3D). In contrast, the Z-score was lowest under conditions with TMAC added (Fig. 3D). We then investigated the effect of TMAC at concentrations above 10 mM. It has been reported that high concentrations of TMAC inhibit PCR [36]. Therefore, we amplified cDNA by adding 25 mM, 50 mM, and 100 mM TMAC and examined the effect on the amount of cDNA (Fig. S8). A clear decrease in byproduct DNA was confirmed under conditions of 25 mM and 50 mM TMAC. As the TMAC concentration increased, the amount of cDNA tended to decrease, and DNA amplification was completely inhibited under conditions of 100 mM TMAC (Fig. S8). Because high concentrations of TMAC were expected to affect the elongation activity of terminal transferase, subsequent experiments were performed using a concentration of 10 mM. The results revealed that adding 10 mM TMAC to the poly-A tailing step specifically inhibited the amplification of byproduct DNA.

In the above experiments, we attempted to stabilize dA/dT base pairs by replacing the substrate of terminal transferase and adding compounds to the buffer solution. Similarly, we hypothesized that adding modifications to the oligo-dT sequence of reverse-transcription primers to enhance dA/dT base pairs would specifically suppress byproduct DNA. The 5-hydroxybutynl-2′-deoxyuridine (Super T) modification has been shown to stabilize dA/dT base pairs [37]. We prepared primers with four Super T modifications in the 24-base oligo-dT region of the reverse-transcription primer. We amplified cDNA using both unmodified and modified primers and compared the results (Fig. S9). This showed that the amount of byproduct DNA decreased and the Z-score also decreased. Additionally, the amount of cDNA under Super T-modified conditions was equivalent to or greater than the average amount of cDNA under unmodified conditions (n=4: 137.4±12%).

Thus, by adding factors that enhance the stability of dA/dT base pairs to the substrate of terminal transferase, reaction solution, and reverse-transcription primers, the amount of byproduct DNA amplification could be reduced without decreasing the amount of amplified cDNA.

### Effects of 2F-dATP and TMAC on the poly-A tailing reaction in relation to the length of the oligo-dT primer

The results of the previous experiment demonstrated that the investigated factors (2F-dATP, TAA, and Super T modification in the primer) that enhance the stability of dA/dT base pairs specifically suppress the amplification of byproducts of poly-A tagging. Next, we investigated the effect of factors stabilizing dA/dT base pairs on the length of the poly-A tail added to the oligo-dT primer. We present a typical example of the effect of factors that stabilize dA/dT base-pairing on the length of the poly-A tail added to the oligo-dT primer in Fig. 4. Under conditions without stabilizing factors, the size distribution of poly-A-tailed oligo-dT primers exhibited a reproducible bimodal pattern in all technical replicates (n=4) (Fig. 4 and Fig. S10). This result was consistent with the results shown in Fig. S2 and Fig. S3. Under conditions with 2F-dATP or TMAC, the very broad peak observed at 48 bases or more disappeared in all technical replicates (n=4) (Fig. 4 and Fig. S10). Meanwhile, only sharp peaks below 48 bases were observed. This indicated that, under conditions with a stabilizing factor, poly-A is added to almost all oligo-dT, but its length is kept short. Next, we applied an FAM modification to the 5′ end of Super T-modified oligo-dT primers and investigated the effect of Super T modification on poly-A tailing (Fig. S11). Under the Super T-modified conditions, the signal of the very broad peak at 48 bases or more was significantly weakened compared with that under the unmodified conditions (Fig. S11). Sharp peaks at 48 bases or less were observed in both conditions. These results indicate that Super T modification of the oligo-dT sequence suppresses poly-A tailing of reverse-transcription primers.

**Figure 4.**
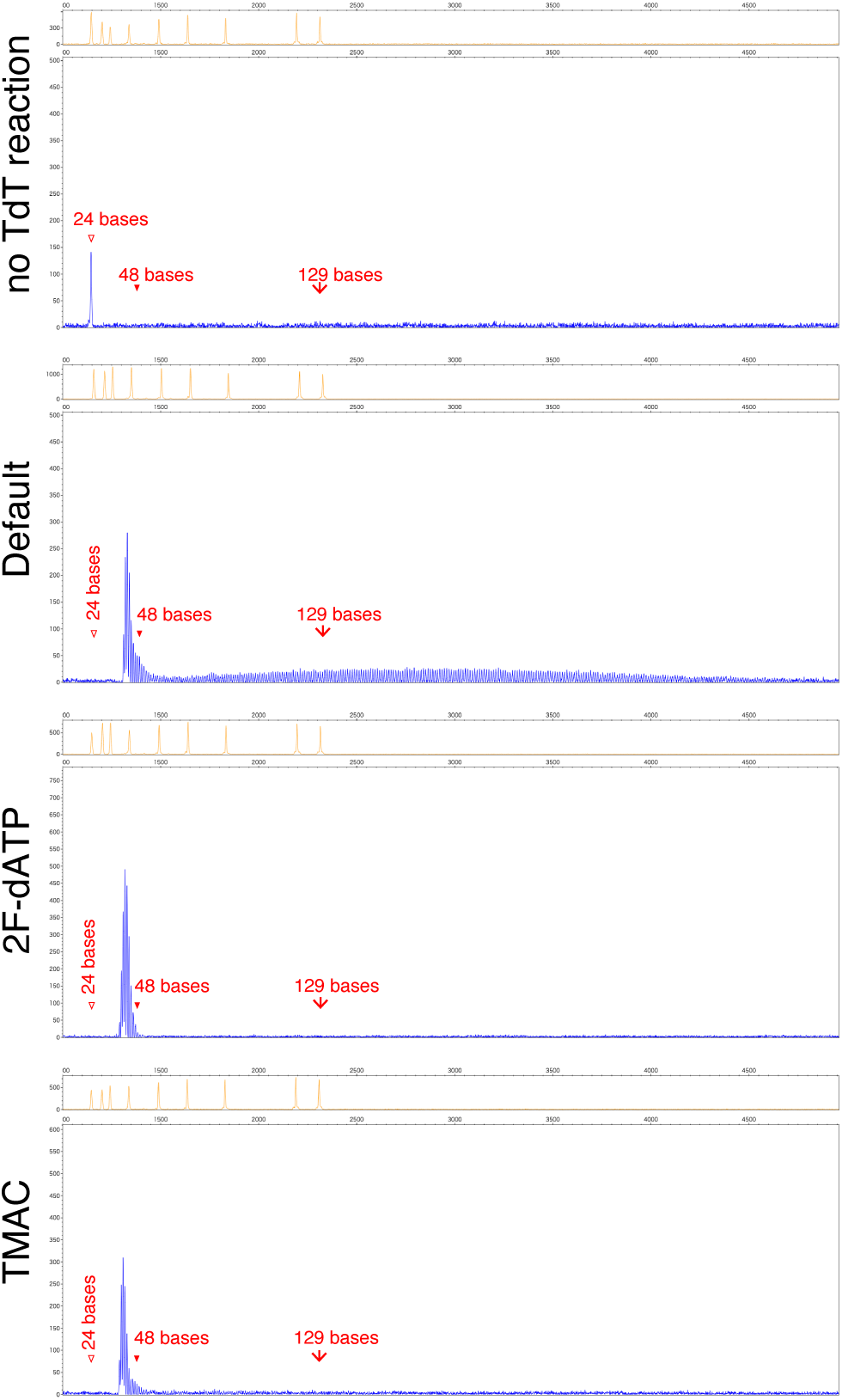
Effects of 2F-dATP and TMAC on the length of poly-A-tailed oligo-dT primers. We performed a 7-min poly-A tailing reaction on FAM-labeled oligo-dT and analyzed the length by capillary electrophoresis. Technical replicates (n=4) were performed, and typical patterns were observed. All electrophoresis patterns are shown in Fig. S10. Under the “2F-dATP” condition, 2F-dATP was used in place of dATP. Under the “TMAC” condition, 10 mM TMAC was added. The “no TdT reaction” condition did not undergo the TdT reaction and is a negative control. Standard DNA was prepared using LIZ120.

Thus, factors known to enhance the stability of dA/dT base pairs, such as modified nucleotides, TAA compounds, and base modification of oligo-dT sequences, suppress poly-A tailing of reverse-transcription primers and, in particular, inhibit the formation of longer poly-A sequences.

### Improvement of usability in whole-transcript amplification by poly-A tagging using SupTag

We successfully suppressed the amplification of byproduct DNA in poly-A tagging by adding a factor that stabilizes dA/dT base pairs in the poly-A tailing reaction. We named this method SupTag. Next, we investigated the impact of SupTag on the usability of WTA using the poly-A tagging method. First, we performed poly-A tagging-based WTA using different adapter sequences to ensure that suppression PCR did not occur, and then examined the effect of SupTag on WTA (see Fig. 5, upper panels). We used a set of conditions that combined 25% 2F-dATP and 10 mM TMAC for SupTag. After WTA, we performed column purification to recover DNA fragments of 70 bp or longer and examined the size distribution of the amplified cDNA. Amplicon DNA of 131 bp in size was observed regardless of the presence or absence of SupTag. This was not observed under suppression PCR conditions (Fig. 5). The 131-bp amplicon DNA was considered to be a dimer of the reverse-transcription primer and the tagging primer. This amplicon DNA was completely removed by one round of bead purification. DNA fragments ranging from 150 bp to 450 bp were byproduct DNA generated by poly-A tagging, and even after three rounds of bead purification, byproduct DNA longer than 300 bp could not be completely removed (Fig. 5, asterisks). Even under conditions with suppression PCR, longer byproduct DNA was observed to remain to some extent even after repeated bead purification (Fig. 5, lower panels). Under SupTag conditions, byproducts were barely observed after column purification. Thus, SupTag suppressed the amplification of byproducts under conditions with or without suppression PCR. This eliminated the need for repeated bead purification, thereby reducing the number of experimental steps.

**Figure 5.**
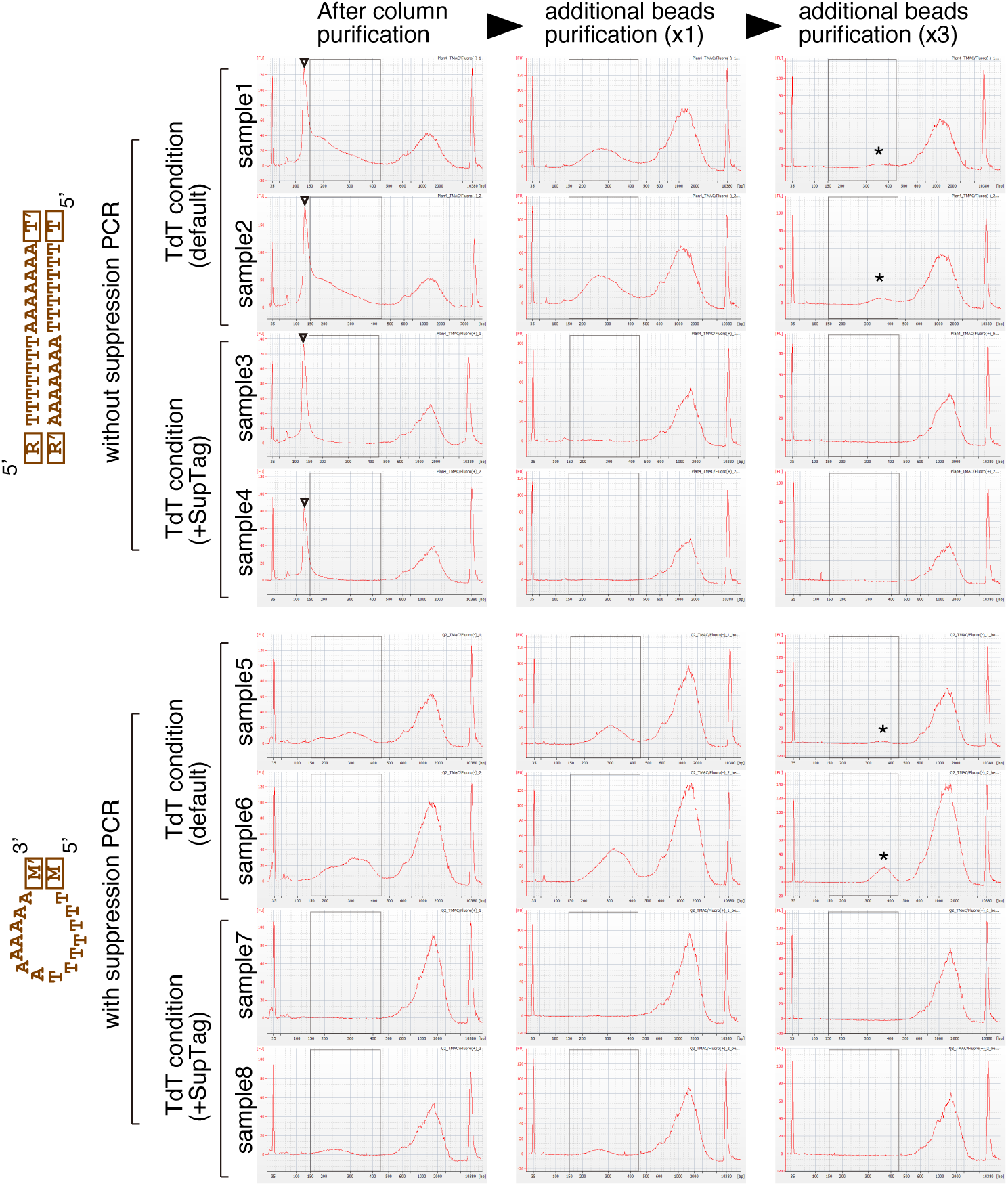
Effects of suppression PCR and SupTag on byproduct DNA. The lower panels show electrophoresis patterns of amplified cDNA obtained using adapter sequences that enable suppression PCR to function during PCR amplification. The upper panels show electrophoresis patterns of amplified cDNA obtained using adapter sequences with different ends during PCR amplification. In the conditions of the upper panels, suppression PCR does not function. WTA by poly-A tagging was performed under normal conditions and SupTag conditions, each with two technical replicates. Under SupTag conditions, 25% of dATP in the poly-A tailing reaction was replaced with 2F-dATP, and 10 mM TMAC was added. The poly-A tailing reaction was performed for 150 s. The left column shows the electrophoresis pattern of DNA purified by a column after PCR. The central and right columns show the electrophoresis patterns after one or three rounds of bead purification. The amplified DNA indicated by the arrowheads observed at approximately 130 bp represents the dimer of the reverse-transcription primer and tagging primer and appears only under conditions without suppression PCR. The black brackets indicate the range of 150–450 bp containing byproduct DNA. The asterisks indicate byproduct DNA observed after bead purification.

### Application of SupTag to the high-throughput RNA-Seq method and its effects

In the above verification, WTA of poly-A tagging was performed using only PCR tubes. Next, we investigated the effect of poly-A tagging by applying SupTag to the Quartz-Seq2 method, which is used as a single-cell RNA-seq method using a 384-well plate, with the addition of 10 pg of total RNA. Under the same conditions, the copy number of reverse-transcription primers per the same amount of RNA is 10 times higher. First, 10 pg of total RNA at the single-cell level was added to each well of the 384-well plate, and then WTA was performed. Extending the TdT reaction time from 75 to 150 s resulted in the production of a significant amount of byproduct DNA (Fig. S12A). Fourteen rounds of bead purification were repeated to completely remove these byproducts, and their removal was confirmed. Because bead purification and DNA electrophoresis for DNA size analysis took approximately 40–50 min per round, a very long experimental time was required. Under SupTag conditions, however, the initial amount of byproduct DNA was low, and nearly all of it could be removed with a single purification step.

Subsequently, we prepared a DNA sequencing library using amplified cDNA before and after byproduct DNA removal under Default conditions, as well as using amplified cDNA from SupTag. These DNA sequencing library samples were mixed at the same molar concentration and analyzed using an Illumina short-read sequencer (NextSeq500). The results showed that the number of reads obtained per sequence run differed by approximately 1.64-fold between the two conditions when the same WTA cDNA was purified once and when it was purified 14 times (Fig. S12C). However, the number of reads from WTA cDNA purified 14 times and WTA cDNA using SupTag differed by less than 4%. In Read2 of the sequence reads, transcript sequences are read, and because byproduct DNA does not contain transcript sequences, it is expected that poly-A or poly-T sequences will be read. Therefore, when analyzing over-represented sequences in the Read2 sequences, homopolymer sequences containing poly-T and poly-A were confirmed. Under the Default conditions, after one round of bead purification, homopolymer sequences accounted for approximately 0.9±0.1% of the total sequences. They were almost absent under the other conditions. The unique mapping ratios under each set of conditions were as follows. Under the Default conditions, after one round of bead purification, the unique mapping ratio was 71.2±0.2% (n=2). Meanwhile, under the Default conditions with 14 rounds of bead purification, it was 73.1±0.2% (n=2). Under the SupTag conditions, a single round of bead purification yielded a unique mapping ratio of 73.2±0.3% (n=2). In conditions marked by a significant yield of byproducts, the distinctive mapping ratio exhibited a modest decline. In the single-bead purification sample under the Default conditions, in which a substantial amount of byproduct DNA was detected, a decrease in unique molecular identifier (UMI) count was also detected. To confirm the reproducibility of this result, we performed the same experiment using NextSeq1000 and MiSeq, which are different models of Illumina sequencers. The results showed a similar trend to that observed with the NextSeq500. Thus, in Illumina short-read sequencers, under conditions where a large amount of byproduct DNA is present, the number of reads obtained per sequence run decreases significantly, and a decrease in the unique mapping ratio is also observed (Fig. S12A and B).

Next, we analyzed samples that did not show a significant decrease in the number of sequence reads in more detail to investigate qualitative changes in amplified cDNA using SupTag. To this end, we quantified gene expression using fastq sequence reads obtained under the Default conditions (14 rounds of purification) and SupTag conditions (1 round of purification), which contained almost no byproduct DNA, with an average data volume of 160,000 reads per CB (Fig. S12B). The results revealed that the average UMI count of 384 wells was approximately 9.3% higher for SupTag (Default beads ×14: 49,066±403 [n=2], SupTag beads ×1: 53,609±735 [n=2]). Additionally, the average gene count of 384 wells was approximately 4.2% higher for SupTag (Default beads ×14: 8,676.8±43.3 [n=2], SupTag beads ×1: 9,045.7±93.7 [n=2]). To provide deeper insight into the efficiency of mRNA capture, the ratio of unique mapped reads to UMI count is referred to as the UMI filtering ratio. A higher value indicates fewer PCR duplicates and higher diversity of cDNA molecules in the DNA sequencing library. The average UMI filtering ratio was approximately 9.6% higher in SupTag (Default beads ×14: 0.456±0.001 [n=2], SupTag beads ×1: 0.5±0.004 [n=2]). Under conditions where a large amount of byproducts are generated, the SupTag conditions, under which there is less amplification of byproduct DNA, showed higher detection sensitivity in the quantification of gene expression. Under these conditions, where large amounts of byproducts are amplified, the quality of cDNA used in the DNA sequencing library deteriorates, which lowers the sensitivity with which gene expression can be quantified. In contrast, SupTag suppresses the amplification of byproduct DNA, suggesting that high quantification performance can be maintained.

In SupTag, factors that stabilize dA/dT base pairs are added to the poly-A tailing reaction using TdT. The effects of these factors on the preference for poly-A addition to cDNA and, consequently, on transcriptomic analysis results remain unclear. To minimize the influence of byproduct DNA and investigate the effect of SupTag on cDNA, we compared conditions with and without SupTag under conditions where byproduct DNA amplification was minimal and such DNA was easily removable. Under the same experimental conditions, we reduced the TdT reaction time to 75 s to decrease the amount of byproduct DNA in the Default conditions (Fig. S13). Furthermore, by performing bead purification three to four times, the byproduct DNA was completely removed. Under SupTag conditions, no amplification of byproduct DNA was observed. The numbers of sequence reads from WTA cDNA (Default) and WTA cDNA using SupTag differed by less than 4%. Sequence analysis using this amplified cDNA showed that there was little difference (<0.4%) between the number of genes detected and the UMI count. Next, clustering analysis was performed without removing the batch effect, and the samples derived from 10 pg of total RNA (1,536 wells) were divided into two clusters. Cluster 0 contained 99.3% SupTag-derived samples, while Cluster 1 contained 98.6% Default-derived samples. Furthermore, analysis of differentially expressed genes between these two clusters identified only 14 genes, including *ACTB* and *RPL41*. Although these genes differed in their expression level, no cases of undetectable gene expression were observed. These results suggest that SupTag has only a very limited effect on the poly-A tailing reaction in cDNA and can suppress byproduct DNA amplification in a highly specific manner.

## Discussion

In this study, we investigated the molecular mechanism of byproduct DNA amplification in the poly-A tagging method and developed a specific method for suppressing it using SupTag. Researchers have been aware of the issue of byproduct DNA associated with poly-A tagging since the report by Iscove et al. published in 2002, but there has been no effective solution to it based on a molecular-level understanding of how byproduct DNA is synthesized [19][20]. We propose SupTag as a potential solution to this issue, a schematic representation of which is shown in Fig. 6. In conventional poly-A tagging, the addition of poly-A by TdT results in the oligo-dT reverse-transcription primer having a long poly-A tail, which facilitates annealing of the tagging primer and amplification of byproduct DNA. In contrast, in SupTag, the dA/dT base pairs formed by poly-A-tailed oligo-dT primers are stabilized, thereby inhibiting further elongation of the poly-A tail. As a result, it was suggested that annealing of the tagging primer becomes difficult, leading to specific suppression of the amplification of byproduct DNA. SupTag makes use of TdT, which significantly reduces the efficiency of base addition to the 3′ recessive end. This enables it to adopt a strategy of inhibiting poly-A addition to oligo-dT primers, which serve as precursors for byproduct DNA [31]. Furthermore, under conditions where large amounts of byproduct DNA are amplified, even if the byproduct DNA is physically removed, the numbers of UMIs and genes detected are lower than in the SupTag conditions, as demonstrated by our results (Fig. S12). This is consistent with previous reports by Iscove et al. and Kurimoto et al., who reported that the detection sensitivity for quantifying gene expression decreases when a large amount of byproduct DNA is amplified [19][20]. It was anticipated that the sequencing library containing byproduct DNA would yield a large number of sequences derived from byproduct-derived poly-A or poly-T, but only about 1% were detected. These results suggest that sequencing libraries derived from byproduct DNA are difficult to read using an Illumina sequencer and reduce the number of fastq reads. Previously, Iscove et al. reported that byproduct DNA reduces transcript-derived signal intensity and detection sensitivity on a microarray platform [19]. Although the detection platforms differ, the results of this study are consistent in that the presence of byproduct DNA reduced the number of reads and the detection sensitivity. Given that numerous other sequencing platforms are available, careful evaluation of the impact of byproduct DNA on their analytical performance should be performed, including for newly developed platforms that are soon to be released [38][39][40].

**Figure 6.**
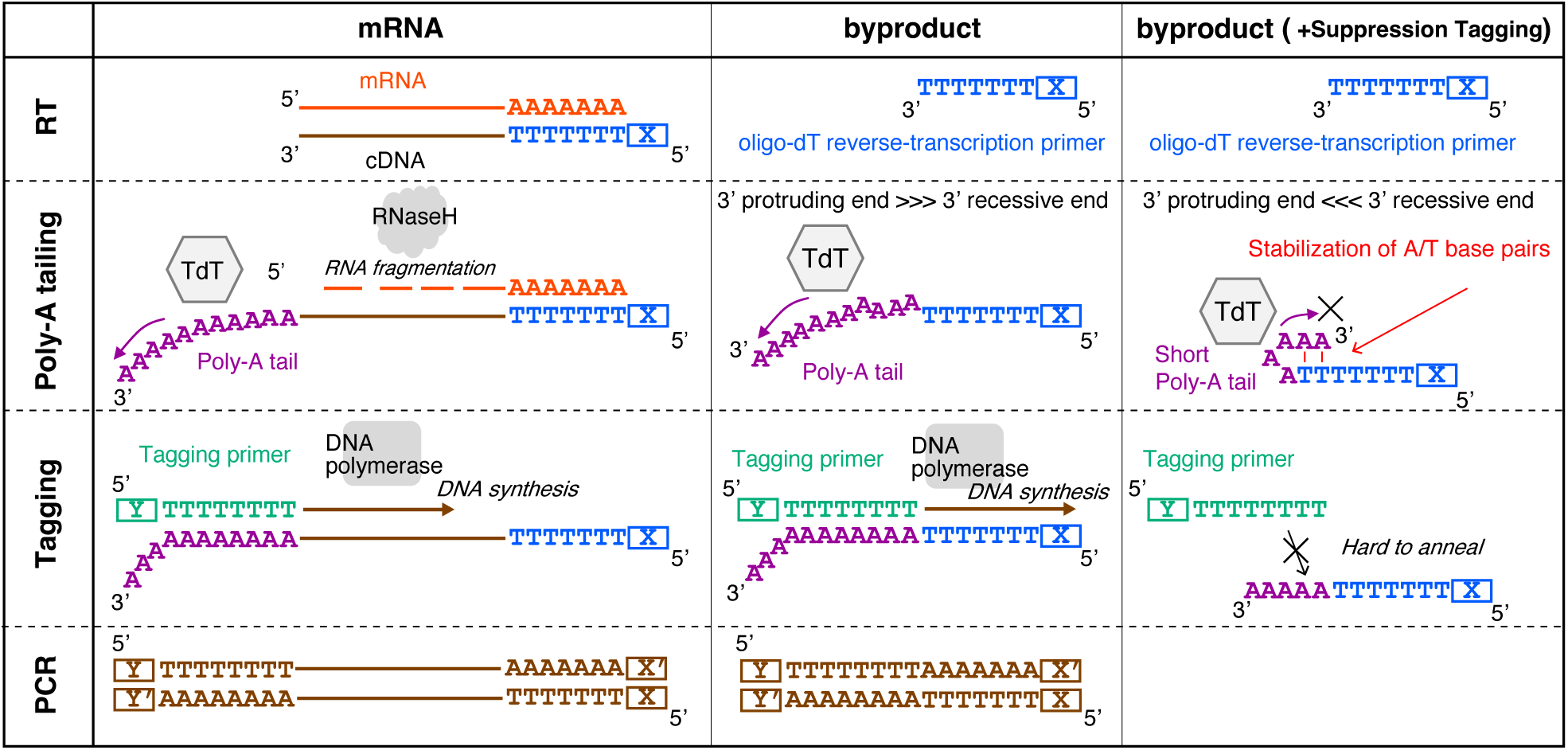
Schematic diagram of SupTag for suppression of amplification of byproduct DNA. The left column shows the mechanism of cDNA amplification by poly-A tagging from mRNA. The central column shows the mechanism of amplification of byproduct DNA under normal conditions. The right column shows a schematic diagram of the mechanism of inhibition of amplification of byproduct DNA by SupTag.

Poly-G tagging and poly-C tagging have also been reported as WTA methods using terminal transferase [41][42][43]. In addition, in a single-cell RNA-seq method using WTA by poly-C tagging and barcode beads, a strategy has been reported to suppress the amplification of byproduct DNA by adding a modified nucleotide called ddCTP, which is a dideoxynucleotide triphosphate, during the TdT reaction [43]. ddCTP is a modified nucleotide that completely stops elongation by DNA polymerase, and when added in low concentrations, it is incorporated equally into byproduct DNA and single-stranded cDNA. Referring to this paper, we performed experiments by adding ddATP to our WTA method using poly-A tagging. In poly-A tailing with ddATP, the probability that extension of a poly-A tail would stop within 10 nucleotides reached 40.1% at 5% ddATP and 94.3% at 25% ddATP. We observed that the cDNA yield decreased to approximately 1/4 at 5% replacement and to approximately 1/80 at 25% replacement (ddATP 5%: 27.2±4.5%, ddATP 25%: 1.2±1.3%, n=2). To confirm reproducibility, we repeated this experiment and confirmed that the cDNA yield decreased to approximately 1/8 at 5% addition and to approximately 1/120 at 25% addition (ddATP 5%: 12.6±6.5%, ddATP 25%: 0.4±0.8%, n=2). These results indicate that the ddNTP addition strategy inhibits cDNA amplification by our poly-A tagging method, which is significantly different from SupTag, which did not impair cDNA yield. Thus, the ddNTP addition strategy was not effective for our amplification method. Additionally, because SupTag is a technology based on the stabilization of dA/dT base pairs, it is effective for poly-A tagging using oligo-dT primers but has technical limitations that prevent its application to methods involving poly-T, poly-C, or poly-G tailing.

Components available for use with SupTag include Super T modification of oligo-dT primers, modified nucleotides, and chemicals (such as TMAC), but their costs and practicality vary significantly. In the case of 5-hydroxybutynl-2′-deoxyuridine (Super T) modification, primers were prepared with four modifications at the 24-base oligo-dT region of the reverse-transcription primer. However, adding four modifications increased the price of the reverse-transcription primer by 12.6 times. Super T-modified primers are extremely expensive and are not cost-effective, especially when a large number of cell barcode-labeled reverse-transcription primers are required. In contrast, the cost of TMAC is extremely low, at only ¥0.005 (US$0.00003) per 384-well plate. Meanwhile, when using 25% 2F-dATP, the additional cost is ¥255 (US$1.66) per 384-well plate. TMAC is also highly practical because it can be easily added to existing reaction systems and can be recommended as a standard condition for SupTag.

The successful development of SupTag has strengthened control over byproduct DNA and enabled reduction of its amount without the need for conventional methods such as suppression PCR. Suppression PCR prevents the amplification of short byproduct DNA by having identical complementary sequences at both ends of cDNA, but it has the limitation that different sequences cannot be added to the two ends [8][26]. SupTag eliminates this limitation and allows the adapter sequences used for reverse-transcription primers to be different from those used for tagging primers (Fig. 5). In high-throughput long-chain cDNA analysis, such as single-cell RNA-seq, it is important that the adapter sequences at the two ends are different [44][45]. For example, when using scNanoGPS, data analysis software for performing high-throughput long-read cDNA analysis, the program needs to accurately recognize adapter sequences adjacent to barcode sequences used to distinguish samples or single cells [45]. For this reason, it is essential that the sequences at the two ends are different and that adjacent adapter sequences can be recognized to identify barcode sequences. The characteristic of having different sequences at the two ends is particularly important in high-throughput long cDNA analysis. SupTag is expected to enable the development of Quartz-Seq2, which has high sensitivity, into a method compatible with long-read analysis. Finally, byproduct DNA generated by poly-A tagging tends to have a greater impact on the analysis as the amount of RNA decreases, making it difficult to accurately quantify gene expression. However, the use of SupTag demonstrates the potential to apply poly-A tagging stably to transcriptomic analysis using even minute RNA samples, which is expected to contribute to improving the accuracy of single-cell analysis and the analysis of low-abundance samples.

## Experimental procedures

### Method of whole-transcript amplification with poly-A tagging

To investigate the effects of various candidate factors on the whole-transcript amplification method using poly-A tagging, the following WTA method was used. Based on previously reported WTA methods using poly-A tagging [8], we modified the reverse-transcription reaction performed in a 384-well plate to a PCR tube so that multiple conditions could be tested simultaneously. Specifically, the following reactions were performed. A total of 10 μL of RT premix (2× Thermopol buffer, 1.25 U/μL of SuperScriptIII, 0.137 U/μL of RNasin Plus) was added to 10 μL of lysis solution (1 ng of total RNA, 0.3% NP-40, 0.12 mM dNTPs, 0.111 μM Q2v3.2RT_Primer, RNasin Plus 1 U/μL), mixed thoroughly, and then reverse transcription was performed. Total RNA derived from human iPS (201B7) or universal mouse reference RNA (UMRR) was used. The reverse-transcription reaction was performed at 35°C for 5 min, followed by 50°C for 50 min. The reverse-transcription reaction solution was purified using Zymo DNA Clean&Concentrator-5. We added 25 μL of TdT premix (1× Thermopol buffer, 2.4 mM dATP, 0.0384 U/μL RNase H, 26.88 units/μL terminal transferase [TdT]) to 20 μL of purified cDNA solution and mixed thoroughly. The mixture was then incubated at 37°C for 75 s or 37°C for 150 s to perform the poly-A tailing reaction. The solution was inactivated by heating it to 65°C for 10 min. When adding modified nucleotides, 25% or 10% of the dATP was replaced with the modified nucleotides. For TAA, 10 mM was added to the TdT premix. Next, the poly-A tagging solution was heated to 98°C for 130 s and then cooled to 40°C for 1 min. Subsequently, the temperature was increased by +0.2°C per cycle and maintained for 1 s, and this cycle was repeated 140 times to raise the temperature to 68°C, after which second-strand synthesis was performed at 68°C for 5 min. Next, we added 50 μL of PCR premix (0.996× Mighty Amp Buffer version 2, 1.895 μM gM_primer_Q2) and mixed well. Subsequently, PCR amplification was performed under the following temperature conditions for 12 cycles: 98°C for 10 s, 65°C for 15 s, and 68°C for 5 min. The mixture was then incubated at 68°C for an additional 5 min. In this amplification, both ends of the amplified cDNA have the same sequence, and suppression PCR works by using a single PCR primer. The PCR solution was purified using a Qiagen MinElute column, followed by additional purification with Ampure XP. The size distribution of the amplified DNA was determined using the Bioanalyzer 2100 High Sensitivity DNA kit. DNA fragments between 100 and 450 bp were quantified as byproduct DNA, while fragments between 550 and 9,000 bp were quantified as amplified cDNA. The ratio of byproduct DNA to amplified cDNA was calculated to determine the Z-score, which was used as an indicator of byproduct DNA suppression for visualization and figures. For conditions without suppression PCR, the primers were modified as follows. The reverse-transcription primer was RTprimer_10×, and the tagging primer was Tagging_10×TSO. For PCR primers, 10×pTSOprimer and 10×pR1 were used, each at a concentration of 0.947 μM. Detailed sequence information for the primers is provided in the supporting information.

### Fragment analysis of poly-A-tailed oligo-dT primer

To investigate the size distribution of poly-A-tailed oligo-dT primers at single-base resolution, the following experimental procedure was performed. An oligo-dT/oligo-dA primer labeled with the fluorescent substance FAM at the 5′ end was prepared, and poly-A or poly-T tailing was performed using terminal transferase. In this experiment, three types of primers (FAMdT24, FAMdA24, and FAMdT24withSuperT) were used. Detailed information on the primers is provided in the supporting information. Poly-A tailing solution (20 mM potassium acetate, 8 mM Tris-acetate [pH 7.9], 4 mM magnesium acetate, 0.4 mM DTT, 0.3 μM FAMdT24 oligo, 14.93 U/μL terminal transferase [TdT], 1.33 mM dATP) was prepared, reacted at 37°C for 1.25 to 7 min, and then inactivated at 65°C for 10 min. Individual reaction times are indicated in the legend of each figure. When using 2F-dATP, it was added at a concentration of 1.33 mM instead of dATP. When adding TAA to the poly-A tailing solution, the final concentration was adjusted to 10 mM. The poly-A tailing solution was diluted with nuclease-free water, supplemented with a size standard (GeneScan 120LIZ Size Standard, Applied Biosystems PN.4324287), and detected using an ABI 3130xl (Thermo Fisher Scientific). Unreacted FAM-oligo-dT was detected as 8.66±0.05 (n=13) bp shorter. Meanwhile, unreacted FAM-oligo-dA was detected as 16.59±0.09 (n=12) bp shorter. The base length corrected for the above detection differences is shown in the figure.

### Melting curve analysis of poly-A-tailed oligo-dT primer

A total of 10 μl of Poly-A tailing solution (20 mM Tris-HCl [pH 8.8], 10 mM (NH4)_2_SO_4_, 10 mM KCl, 2 mM MgSO_4_, 0.1% Triton X-100, 0.3 μM dT24 oligo, 14.93 U/μL Terminal Transferase, 1.33 mM dATP, 1× Rox dye, 1× EvaGreen dye) was prepared. The poly-A tailing solution was heated at 37°C for 75 s, inactivated at 95°C for 15 s, and then cooled to 4°C at a rate of −1.6°C/s. Then, it was heated to 95°C at a rate of +0.05°C/s, and the fluorescence intensity was measured during the heating process. For poly-T tailing, dTTP was used instead of dATP. Additionally, when using an oligo-dA primer for the template DNA, dA24 was used instead of dT24. Detailed primer sequences are provided in the supporting information.

### Whole-transcript amplification method using poly-A tagging and preparation of DNA sequencing library for high-throughput RNA-seq using 10 pg of total RNA

To evaluate the WTA method with poly-A tagging at the single-cell level, 10 pg of purified total RNA was analyzed. Total RNA derived from human iPS (201B7) was kindly provided by Ms. Yasuko Hisano and Dr. Takeo Yoshikawa of the Laboratory for Molecular Psychiatry, RIKEN Center for Brain Science [46]. Poly-A tagging-based WTA and DNA sequencing library preparation using a 384-well plate were performed in accordance with the method previously reported in our Quartz-Seq2 paper [8]. We used the Low Volume Chip of the MANTIS Liquid Dispenser (Formulatrix) to dispense 0.1 μl of purified 100 pg/μl human iPS cell (201B7) total RNA into all wells of a 384-well lysis plate. This resulted in 10 pg of total RNA being dispensed per well. Under normal conditions, the poly-A tailing reaction is performed for 75 s, but we also prepared conditions where it was intentionally performed for 150 s. Under SupTag conditions, 25% of dATP in the poly-A tailing reaction was replaced with 2F-dATP, and 10 mM TMAC was added. The reaction time for poly-A tailing is indicated in the legend of the figure. The RT primer used for the 384-well plate was the v3.2a type. DNA sequencing library was prepared using amplified DNA, and DNA quantification was performed using the Bioanalyzer 2100 High Sensitivity DNA kit and Qubit Flex. The quantification values from both methods showed a very high positive correlation (y=1.0249x, R^2^=0.9961, n=6).

### Analysis of high-throughput RNA sequencing data

Each DNA sequencing library was mixed at an equal molar concentration and analyzed using an Illumina sequencer. Upon analysis with a NextSeq1000 sequencer, after mixing the DNA sequencing library, PhiX Control v3 was mixed at an equimolar concentration with the mixture, and sequence analysis was performed. The read lengths for sequencing analysis using the NextSeq 500/550 High Output v2 Kit (75 cycles) were as follows (Read1, 23 cycles; Index1, 6 cycles; Read2, 63 cycles). The read lengths for sequence analysis using the NextSeq 1000/2000 P2 Reagents (100 Cycles) v3 were as follows (Read1, 26 cycles; Index1, 6 cycles; Read2, 64 cycles). The read lengths in the sequence analysis using the MiSeq Reagent Nano Kit v2 were as follows (Read1, 23 cycles; Index1, 6 cycles; Read2, 101 cycles). We used the Quartz-Seq2 pipeline (https://github.com/rikenbit/Q2-pipeline_v2) to calculate and generate a digital expression matrix from the bcl files. In the Quartz-Seq2 pipeline, we used bases 1–23 of the Read1 fastq file as the cell barcode sequence and the molecular barcode sequence, and bases 1–62 of the Read2 fastq file as the transcript sequence. We used the GRCh38 primary genome assembly as the reference genome and the Gencode Release 44 (GRCh38) gtf file for gene annotation.

We enabled the option to include mapping reads within introns in the count and quantified the gene expression. To analyze the gene expression matrix and visualize the results, we used Seurat 4.1.1 and performed data analysis using the standard procedure described in the tutorial [47].

## Data availability

The raw sequencing data and the processed digital expression matrix in this study have been deposited in the Gene Expression Omnibus (GEO) database under accession number GSE303412. The software code for the Quartz-Seq2 pipeline is available at https://github.com/rikenbit/Q2-pipeline_v2 (DOI: 10.5281/zenodo.15920498).

## Supporting information

This article contains supporting information. In addition, the following reference is cited in this supporting information.

Kim, S., Chen, J., Cheng, T., Gindulyte, A., He, J., He, S. et al. (2025) PubChem 2025 update *Nucleic Acids Res* 53, D1516-D1525

## Acknowledgments

We would like to thank the members of the Omics AI Research Team at RIKEN, Akihiro Matsushima and Takumi Ichikawa, for their management of the IT infrastructure. We are also grateful to the Support Unit for Bio-Material Analysis, RIKEN CBS Research Resources Division, for technical help with the analysis of single-stranded DNA sizes. Moreover, we would like to express our gratitude to Ms. Yasuko Hisano and Dr. Takeo Yoshikawa of the Laboratory for Molecular Psychiatry, RIKEN Center for Brain Science, for providing the total RNA. Finally, we are grateful for English editing support from Edanz (https://jp.edanz.com/ac). This service was provided solely for language editing and did not influence the study design, data collection, analysis, or conclusions.

## Author contributions

Y.S. and I.N. designed the study; Y.S. and Y.I. performed the experiments and analyzed data; M.Y. supported sequence analysis; Y.S. and I.N. wrote the manuscript; I.N. supervised the project.

## Funding and additional information

This research was supported by JST K Program Grant Number JPMJKP23H5, AMED under Grant Number JP21bm0404073, and JSPS KAKENHI Grant Number JP19H03214. Sequencing analyses were supported by the Medical Research Center Initiative for High Depth Omics program.

## Abbreviations and nomenclature

RNA sequencing: RNA-seq Suppression
Tagging: SupTag
WTA: whole-transcript amplification
bp: base pair
FAM: fluorescein
RT: reverse transcription
PCR: polymerase chain reaction
mRNA: messenger RNA
IVT: in vitro transcription
TdT: terminal transferase / terminal deoxynucleotidyl transferase
dA: 2′-deoxyadenosine
dT: 2′-deoxythymidine
dNTP: deoxyribonucleotide triphosphate
dATP: 2′-deoxyadenosine 5′-triphosphate
dTTP: 2′-deoxythymidine 5′-triphosphate
dZTP: 2-amino-2’-deoxyadenosine 5′-triphosphate
NH2dATP: 2′-amino-2′-deoxyadenosine 5′-triphosphate
AzdATP: 2′-azido-2′-deoxyadenosine 5′-triphosphate
OmdATP: 2′-O-methyladenosine 5′-triphosphate
2F-dATP: 2′-fluoro-2′-deoxyadenosine triphosphate
TAA: tetraalkylammonium chloride
TMAC: tetramethylammonium chloride
TEAC: tetraethylammonium chloride
TPrAC: tetrapropylammonium chloride
TBAC: tetrabutylammonium chloride
TPeAC: tetrapentylammonium chloride
ddCTP: 2′,3′-dideoxycytidine-5′-O-triphosphate
ddNTP: dideoxynucleotide triphosphate
ddATP: 2′,3′-dideoxyadenosine-5′-O-triphosphate
UMI: unique molecular identifier

## References

[1]. Stark, R., Grzelak, M., and Hadfield, J. (2019) RNA sequencing: the teenage years Nat Rev Genet 20, 631–656

[2]. Huang, K., Xu, Y., Feng, T., Lan, H., Ling, F., Xiang, H., et al. (2024) The Advancement and Application of the Single-Cell Transcriptome in Biological and Medical Research Biology (Basel) 13,

[3]. Islam, S., Kjällquist, U., Moliner, A., Zajac, P., Fan, J. B., Lönnerberg, P. et al. (2011) Characterization of the single-cell transcriptional landscape by highly multiplex RNA-seq Genome Res 21, 1160–1167

[4]. Hagemann-Jensen, M., Ziegenhain, C., and Sandberg, R. (2022) Scalable single-cell RNA sequencing from full transcripts with Smart-seq3xpress Nat Biotechnol 40, 1452–1457

[5]. Hahaut, V., Pavlinic, D., Carbone, W., Schuierer, S., Balmer, P., Quinodoz, M. et al. (2022) Fast and highly sensitive full-length single-cell RNA sequencing using FLASH-seq Nat Biotechnol 40, 1447–1451

[6] Zheng, G. X., Terry, J. M., Belgrader, P., Ryvkin, P., Bent, Z. W., Wilson, R., et al. (2017) Massively parallel digital transcriptional profiling of single cells Nat Commun 8, 14049

[7]. Rodriques, S. G., Stickels, R. R., Goeva, A., Martin, C. A., Murray, E., Vanderburg, C. R. et al. (2019) Slide-seq: A scalable technology for measuring genome-wide expression at high spatial resolution Science 363, 1463–1467

[8]. Sasagawa, Y., Danno, H., Takada, H., Ebisawa, M., Tanaka, K., Hayashi, T., et al. (2018) Quartz-Seq2: a high-throughput single-cell RNA-sequencing method that effectively uses limited sequence reads Genome Biol 19, 29

[9]. Hashimshony, T., Senderovich, N., Avital, G., Klochendler, A., de Leeuw, Y., Anavy, L., et al. (2016) CEL-Seq2: sensitive highly-multiplexed single-cell RNA-Seq Genome Biol 17, 77

[10]. Salmen, F., De Jonghe, J., Kaminski, T. S., Alemany, A., Parada, G. E., Verity-Legg, J. et al. (2022) High-throughput total RNA sequencing in single cells using VASA-seq Nat Biotechnol 40, 1780–1793

[11]. Hughes, T. K., Wadsworth, M. H., Gierahn, T. M., Do, T., Weiss, D., Andrade, P. R. et al. (2020) Second-Strand Synthesis-Based Massively Parallel scRNA-Seq Reveals Cellular States and Molecular Features of Human Inflammatory Skin Pathologies Immunity 53, 878–894.e877

[12]. Poovathingal, S., Davie, K., Borm, L. E., Vandepoel, R., Poulvellarie, N., Verfaillie, A. et al. (2024) Nova-ST: Nano-patterned ultra-dense platform for spatial transcriptomics Cell Rep Methods 4, 100831

[13]. Sasagawa, Y., Hayashi, T., and Nikaido, I. (2019) Strategies for Converting RNA to Amplifiable cDNA for Single-Cell RNA Sequencing Methods Adv Exp Med Biol 1129, 1–17

[14]. Kapteyn, J., He, R., McDowell, E. T., and Gang, D. R. (2010) Incorporation of non-natural nucleotides into template-switching oligonucleotides reduces background and improves cDNA synthesis from very small RNA samples BMC Genomics 11, 413

[15]. Turchinovich, A., Surowy, H., Serva, A., Zapatka, M., Lichter, P., and Burwinkel, B. (2014) Capture and Amplification by Tailing and Switching (CATS). An ultrasensitive ligation-independent method for generation of DNA libraries for deep sequencing from picogram amounts of DNA and RNA RNA Biol 11, 817–828

[16]. Hardwick, S. A., Hu, W., Joglekar, A., Fan, L., Collier, P. G., Foord, C. et al. (2022) Single-nuclei isoform RNA sequencing unlocks barcoded exon connectivity in frozen brain tissue Nat Biotechnol 40, 1082–1092

[17]. Gholamalipour, Y., Karunanayake Mudiyanselage, A., and Martin, C. T. (2018) 3’ end additions by T7 RNA polymerase are RNA self-templated, distributive and diverse in character-RNA-Seq analyses Nucleic Acids Res 46, 9253–9263

[18]. Mu, X., Greenwald, E., Ahmad, S., and Hur, S. (2018) An origin of the immunogenicity of in vitro transcribed RNA Nucleic Acids Res 46, 5239–5249

[19]. Iscove, N. N., Barbara, M., Gu, M., Gibson, M., Modi, C., and Winegarden, N. (2002) Representation is faithfully preserved in global cDNA amplified exponentially from sub-picogram quantities of mRNA Nat Biotechnol 20, 940–943

[20]. Kurimoto, K., Yabuta, Y., Ohinata, Y., Ono, Y., Uno, K. D., Yamada, R. G., et al. (2006) An improved single-cell cDNA amplification method for efficient high-density oligonucleotide microarray analysis Nucleic Acids Res 34, e42

[21]. Verwilt, J., Mestdagh, P., and Vandesompele, J. (2023) Artifacts and biases of the reverse transcription reaction in RNA sequencing RNA 29, 889–897

[22]. Frohman, M. A., Dush, M. K., and Martin, G. R. (1988) Rapid production of full-length cDNAs from rare transcripts: amplification using a single gene-specific oligonucleotide primer Proc Natl Acad Sci U S A 85, 8998–9002

[23]. Ohara, O., Dorit, R. L., and Gilbert, W. (1989) One-sided polymerase chain reaction: the amplification of cDNA Proc Natl Acad Sci U S A 86, 5673–5677

[24]. Tang, F., Barbacioru, C., Wang, Y., Nordman, E., Lee, C., Xu, N. et al. (2009) mRNA-Seq whole-transcriptome analysis of a single cell Nat Methods 6, 377–382

[25]. Mickelsen, S., Snyder, C., Trujillo, K., Bogue, M., Roth, D. B., and Meek, K. (1999) Modulation of terminal deoxynucleotidyltransferase activity by the DNA-dependent protein kinase J Immunol 163, 834–843

[26]. Sasagawa, Y., Nikaido, I., Hayashi, T., Danno, H., Uno, K. D., Imai, T., et al. (2013) Quartz-Seq: a highly reproducible and sensitive single-cell RNA sequencing method, reveals non-genetic gene-expression heterogeneity Genome Biol 14, R31

[27]. Mereu, E., Lafzi, A., Moutinho, C., Ziegenhain, C., McCarthy, D. J., Álvarez-Varela, A. et al. (2020) Benchmarking single-cell RNA-sequencing protocols for cell atlas projects Nat Biotechnol 38, 747–755

[28]. Tang, F., Lao, K., and Surani, M. A. (2011) Development and applications of single-cell transcriptome analysis Nat Methods 8, S6-11

[29]. Kurimoto, K., Yabuta, Y., Ohinata, Y., and Saitou, M. (2007) Global single-cell cDNA amplification to provide a template for representative high-density oligonucleotide microarray analysis Nat Protoc 2, 739–752

[30]. Kuznetsov, S. V., Ren, C. C., Woodson, S. A., and Ansari, A. (2008) Loop dependence of the stability and dynamics of nucleic acid hairpins Nucleic Acids Res 36, 1098–1112

[31]. Deng, G., and Wu, R. (1981) An improved procedure for utilizing terminal transferase to add homopolymers to the 3’ termini of DNA Nucleic Acids Res 9, 4173–4188

[32]. Nicholson, A. L., and Pasquinelli, A. E. (2019) Tales of Detailed Poly(A) Tails Trends Cell Biol 29, 191–200

[33]. Patra, A., Paolillo, M., Charisse, K., Manoharan, M., Rozners, E., and Egli, M. (2012) 2’-Fluoro RNA shows increased Watson-Crick H-bonding strength and stacking relative to RNA: evidence from NMR and thermodynamic data Angew Chem Int Ed Engl 51, 11863–11866

[34]. Gao, J., Nutan, B., Gargouri, D., Pisal, N. D., Do, V., Zubair, M., Alanzi, H., Wang, H., Lee, D., Joshi, N. and Ullah, A. (2024) Unlocking the potential of chemically modified nucleic acid therapeutics Advanced Therapeutics 7, 11, 2400231–2400259

[35]. Melchior, W. B., and Von Hippel, P. H. (1973) Alteration of the relative stability of dA-dT and dG-dC base pairs in DNA Proc Natl Acad Sci U S A 70, 298–302

[36]. Chevet, E., Lemaître, G., and Katinka, M. D. (1995) Low concentrations of tetramethylammonium chloride increase yield and specificity of PCR Nucleic Acids Res 23, 3343–3344

[37]. Belousov, Y. S., Welch, R. A., Sanders, S., Mills, A., Kulchenko, A., Dempcy, R. et al. (2004) Single nucleotide polymorphism genotyping by two colour melting curve analysis using the MGB Eclipse Probe System in challenging sequence environment Hum Genomics 1, 209–217

[38]. Deamer, D., Akeson, M., and Branton, D. (2016) Three decades of nanopore sequencing Nat Biotechnol 34, 518–524

[39]. [preprint] Kokoris, M., McRuer, R., Nabavi, M., Jacobs, A., Prindle, M., Cech, C., et al. (2025) Sequencing by Expansion (SBX) - a novel, high-throughput single-molecule sequencing technology bioRxiv 10.1101/2025.02.19.639056

[40]. Qin, Y., Koehler, S. A., Ling, Y., Yu, S., Zhang, Y., Luo, J., et al. (2025) Fast, cost-effective and flexible DNA sequencing by roll-to-roll fluidics Nat Methods

[41]. Loh, E. Y., Elliott, J. F., Cwirla, S., Lanier, L. L., and Davis, M. M. (1989) Polymerase chain reaction with single-sided specificity: analysis of T cell receptor delta chain Science 243, 217–220

[42]. Belyavsky, A., Vinogradova, T., and Rajewsky, K. (1989) PCR-based cDNA library construction: general cDNA libraries at the level of a few cells Nucleic Acids Res 17, 2919–2932

[43]. Shichino, S., Ueha, S., Hashimoto, S., Ogawa, T., Aoki, H., Wu, B., et al. (2022) TAS-Seq is a robust and sensitive amplification method for bead-based scRNA-seq Commun Biol 5, 602

[44]. Chen, Y., Sim, A., Wan, Y. K., Yeo, K., Lee, J. J. X., Ling, M. H. et al. (2023) Context-aware transcript quantification from long-read RNA-seq data with Bambu Nat Methods 20, 1187–1195

[45]. Shiau, C. K., Lu, L., Kieser, R., Fukumura, K., Pan, T., Lin, H. Y. et al. (2023) High throughput single cell long-read sequencing analyses of same-cell genotypes and phenotypes in human tumors Nat Commun 14, 4124

[46]. Toyoshima, M., Akamatsu, W., Okada, Y., Ohnishi, T., Balan, S., Hisano, Y., et al. (2016) Analysis of induced pluripotent stem cells carrying 22q11.2 deletion Transl Psychiatry 6, e934

[47]. Hao, Y., Hao, S., Andersen-Nissen, E., Mauck, W. M., Zheng, S., Butler, A., et al. (2021) Integrated analysis of multimodal single-cell data Cell 184, 3573-3587.e3529

