## Supporting information for "Elucidation of the mechanism of byproduct DNA synthesis in whole-transcript amplification by poly-A tagging and development of its specific suppression"

### Title

### Authors and affiliations

Yohei Sasagawa (笹川 洋平)[1,2,†], Yoshimi Iwayama (岩山 佳美)[1,2], Minoru Yano (矢野 実)[1], Itoshi Nikaido (二階堂 愛)[1,2,†]

[1]: Department of Functional Genome Informatics, Division of Biological Data Science, Medical Research Laboratory, Institute of Integrated Research, Institute of Science Tokyo, 1-5-45 Yushima, Bunkyo-ku, Tokyo 113-8510, Japan

[2]: Omics AI Research Team, Advanced General Intelligence in Science Program (AGIS), TRIP Headquarters, RIKEN, 2-1 Hirosawa, Wako, Saitama 351-0198, Japan

### Supporting information

**Additional information about experimental procedures**

The table below shows the primer sequences and base modifications used in this study. Notations enclosed in slashes indicate modifiers or modified bases.

| Primer name | Sequence of primer (5' to 3' direction) | Purification |
|---|---|---|
| Q2v3.2RT_Primer | TATAGAATTCGCGGCCGCTCGCGATACNNNNNNNNNNNNNNNNNNNNNNNTTTTTTTTTTTTTTTTTTTTTTTT | HPLC |
| Tagging_primer_Q2 | TATAGAATTCGCGGCCGCTCGCGATTTTTTTTTTTTTTTTTTTTTTTT | HPLC |
| gM_primer_Q2 | GTATAGAATTCGCGGCCGCTCGCGAT | HPLC |
| Q2v3.2RT_PrimerWithSuperT | TATAGAATTCGCGGCCGCTCGCGATACNNNNNNNNNNNNNNNNNNNNNNNT/iSuper-dT/TTTTT/iSuper-dT/TTTTT/iSuper-dT/TTTTT/iSuper-dT/TTTT | HPLC |
| RTprimer_10x | CTACACGACGCTCTTCCGATCTNNNNNNNNNNNNNNNNNNNNNNNNNNNNTTTTTTTTTTTTTTTTTTTTTTTTTTTTTTVN | HPLC |

| | | |
|---|---|---|
| Tagging_10xTSO | AAGCAGTGGTATCAACGCAGAGTACATTTTTTTTTTTTTTTTTTTTT TTT | HPLC |
| 10xpTSOprimer | AAGCAGTGGTATCAACGCAGAG | HPLC |
| 10xpR1 | CTACACGACGCTCTTCCGATCT | HPLC |
| FAMdT24 | /56-FAM/TTTTTTTTTTTTTTTTTTTTTTTT | HPLC |
| FAMdA24 | /56-FAM/AAAAAAAAAAAAAAAAAAAAAAAA | HPLC |
| FAMdT24withSuperT | /56-FAM/T/iSuper-dT/TTTTT/iSuper-dT/TTTTT/iSuper-dT/TTTTT/iSuper-dT/TTTT | HPLC |
| dT24 | TTTTTTTTTTTTTTTTTTTTTTTT | HPLC |
| dA24 | AAAAAAAAAAAAAAAAAAAAAAAA | HPLC |

**Legend of Supplementary Figures**

We have prepared the following legends for each supplementary figure.

**<u>Figure S1</u>**

**Typical pattern of DNA size distribution in whole-transcript amplification by the poly-A tagging method**. Using 1 ng of total RNA derived from human iPS cells, poly-A tailing was performed using the poly-A tagging method with terminal deoxynucleotidyl transferase (TdT) enzyme for 75 s and 150 s. Additionally, conditions were prepared where TMAC and 2F-dATP were added during the poly-A tailing reaction, and under these conditions, byproducts derived from poly-A tagging were hardly observed. The size distribution of the amplified DNA was analyzed using a Bioanalyzer 2100. The x-axis of the figure corresponds to DNA size, measured in base pairs. The y-axis denotes the quantity of fluorescence associated with the DNA. a) shows a typical schematic diagram of amplified cDNA derived from amplified mRNA and byproduct DNA amplified by poly-A tagging. Blue indicates amplified byproduct DNA derived from reverse-transcription primers by poly-A tagging. Red indicates amplified cDNA. b) and c) show the individual electrophoresis patterns and merged patterns under each condition. Under normal conditions for 75 s, the size distribution of byproduct DNA was approximately 100 bp to 300 bp. Under normal conditions for 150 s, the size distribution of byproduct DNA was approximately 100 bp to over 500 bp. The amplified cDNA showed a size distribution ranging from approximately 450 bp to 9000 bp.

**<u>Figure S2</u>**

Analysis of changes in the size of single-stranded DNA from an oligo-dT primer due to non-template-dependent base addition in the TdT reaction. A TdT reaction was performed for 7 min using dATP or dTTP with an oligo-dT primer with FAM attached to the 5′ end, and fragment analysis was performed. Orange indicates detection of the LIZ120 size standard, and blue indicates detection of the FAM fluorescence wavelength. Technical replicates (n=4) were set up, and the third data point from the top was used in Figure 2. “No TdT reaction” refers to a negative control where no terminal transferase reaction was performed. “None-dNTP” indicates a reaction with terminal transferase without adding dNTPs. “+dATP” and “+dTTP” indicate reactions with dATP and dTTP, respectively, followed by a terminal transferase-mediated non-template-dependent base addition reaction.

**Figure S3**

**Changes in the size of single-stranded DNA from oligo-dT primers due to differences in the reaction time of poly-A tailing.** Poly-A tailing reactions were performed for 3 min, 4 min, 5 min, and 7 min, followed by fragment analysis (n=4). LIZ120 was used as the size standard. Typical patterns are shown in the figure. The left panel shows the overall pattern, and the right panel shows an enlarged view.

**Figure S4**

Measurement of the size lengths of FAM-labeled oligo-dT primers and oligo-dA primers using poly-A tailing or poly-T tailing. FAM-labeled oligo-dT primers and oligo-dA primers were prepared, and poly-A or poly-T was added using terminal transferase for 3 min. The size length distribution was analyzed. In each sample, the base with the highest fluorescence intensity was designated as the top peak, and an arrow was placed at its position to indicate the base length. Four technical replicates (n=4) were prepared, and the size distribution was measured and detected. Standard DNA was prepared using LIZ120.

**Figure S5**

**Effect of modified nucleotides on whole-transcript amplification by poly-A tagging.** In poly-A tailing, modified nucleotides were replaced with 25% dATP and WTA was performed (technical replicates, n=3). a) shows the structure of each nucleotide downloaded from PubChem [S1]. b) Z-scores under each condition are shown. c) Electrophoresis patterns of amplified cDNA under each condition are shown.

[S1] Kim, S., Chen, J., Cheng, T., Gindulyte, A., He, J., He, S. et al. (2025) PubChem 2025 update *Nucleic Acids Res* 53, D1516-D1525

**Figure S6**

**Repeatability experiment to investigate the effect of modified nucleotides on WTA by poly-A tagging.** In poly-A tailing, modified nucleotides were replaced with 25% dATP and WTA was performed (technical replicates, n=3). For 2F-dATP, experiments were conducted by replacing 10% of the dATP with 2F-dATP. Electrophoresis patterns of amplified cDNA under each condition are shown. Figure 3 uses electrophoresis images from the central column for dATP and the central column for 2F-dATP (25%).

**Figure S7**

**Effect of poly-A tagging with tetraalkylammonium chlorides (TAA) on WTA**

The reaction time for poly-A tailing using terminal deoxynucleotidyl transferase (TdT) enzyme was set to 150 s. Additionally, each TAA compound was added to the poly-A tailing solution at a concentration of 10 mM. The left two panels show the electrophoresis patterns of amplified cDNA. The right panel shows the structures of TAA downloaded from PubChem [S1].

**Figure S8**

**Effect of TMAC concentration on WTA by poly-A tagging**

TMAC was added to poly-A tailing at concentrations ranging from 20 mM to 100 mM, and WTA was performed. The results of electrophoresis analysis are shown below. b) and c) show the box plots of Z-scores. Because no DNA amplification was detected at a TMAC concentration of 100 mM, the data were excluded from the Z-score display.

**Figure S9**

**The effect of Super T modification of oligo-dT sequences on WTA via poly-A tagging**

a) We show the sequences and base modifications of the reverse-transcription primers used in this experiment. The T in red indicates Super T (5-hydroxybutynl-2′-deoxyuridine) modification. In the “+Super T” condition, reverse-transcription primers containing the Super T modification were used. b) In this experiment, 10 pmol of reverse-transcription primers was used. Poly-A tagging-mediated WTA was performed using different reverse-transcription primers. The poly-A tailing reaction was performed for 2.5 min. The amplified cDNA was subjected to electrophoresis analysis, and the byproduct DNA and amplified cDNA were quantified. c) The Z-score of the ratio of byproduct DNA to amplified cDNA is shown.

**Figure S10**

**Effects of 2F-dATP and TMAC on the length of poly-A-tailed oligo-dT primers**

We performed a 7-min poly-A tailing reaction on FAM-labeled oligo-dT and analyzed the length by capillary electrophoresis. Technical replicates were performed with n=4. Under the “2F-dATP” condition, 2F-dATP was used in place of dATP. In the “TMAC” condition, 10 mM TMAC was added. The “no TdT reaction” condition served as a negative control. Standard DNA was prepared using LIZ120.

**Figure S11**

**Effect of Super T modification on the length of poly-A-tailed oligo-dT primers**

We performed a poly-A tailing reaction for 7 min on a 24-base oligo-dT primer labeled with FAM at the 5′ end and analyzed its length by capillary electrophoresis. The size standard DNA used was LIZ500. Additionally, primers with Super T modifications at four positions in the oligo-dT sequence were prepared and treated with the poly-A tailing reaction. The size of the unreacted 24-base oligo-dT sequence, corrected using the LIZ500 size standard, was 25.67±0.21 bases (n=2) without Super T modification and 28.63±0.14 bases (n=2) with Super T modification. Therefore, for each condition, the size was indicated at positions approximately 1.5 bases and 4.5 bases longer than the size standard DNA.

**Figure S12**

**Effect of SupTag on single-cell RNA-seq under conditions with amplification of large amounts of byproduct DNA**

In this experiment, 10 pg of total RNA (n=384) was amplified into cDNA and gene expression was quantified by the Quartz-Seq2 method. The reaction time for poly-A tailing was 150 s. For each condition, two 384-well plates were analyzed.

a) Electrophoresis patterns of amplified DNA are shown. After column purification, bead purification and electrophoresis analysis were repeated until byproduct DNA was removed, and the electrophoresis patterns are shown. The number of purification steps is indicated within the panel. The panel enclosed in the blue frame shows the electrophoresis patterns of amplified DNA used for sequencing analysis. b) The number of reads was adjusted to 61.44 million reads per 384-well plate, and a digital expression matrix was calculated, the results of which are shown. The input data amount was 160,000 fastq reads per 1 well–10 pg total RNA on average. The UMI filtering ratio was obtained by dividing the number of mapped reads by the number of UMI counts. c) Number of fastq reads obtained per run by each sequencer. d) UMI counts obtained per run by each sequencer.

**Figure S13**

**Effect of SupTag on single-cell RNA-seq under conditions with low levels of amplified byproduct DNA**

We performed the same experiment as in Figure S12, but with a poly-A tailing reaction time of 75 s. For each condition, we analyzed two 384-well plates. a) Electrophoresis patterns of amplified DNA are shown. Under normal conditions, byproducts were completely removed after 3–4 rounds of bead purification. The amplified cDNA shown in the lower panels was used for sequencing analysis. b) Sequencing analysis results showed the following numbers of fastq reads per run (Default-lot1: 39.4M reads, Default-lot2: 39.7M reads, SupTag-lot1: 36.7M reads, SupTag-lot2: 41.5M reads). We calculated the digital expression matrix for each 384-well plate based on the read counts shown below and presented the results. The average read counts per well were 5k, 10k, 20k, 30k, 40k, 50k, 60k, 70k, 80k, and 90k. The Y-axis shows the number of detected genes and the average UMI count. The UMI conversion efficiency is shown for each 384-well plate. c) We used a dataset with an average read count of 90k per well and removed wells with fewer than 2,500 genes detected. The number of wells in each sample after removal is as follows (Default-lot1: 384 wells, Default-lot2: 384 wells, SupTag-lot1: 382 wells, SupTag-lot2: 383 wells). A violin plot of the number of detected genes is shown, separated by 384-well plate conditions and cluster numbers. d) We showed the gene expression levels of all differentially expressed genes in clusters 0 and 1 as Z-scores. A color closer to yellow indicates a higher expression level. e) As an example, the magnitude of gene expression levels of two differentially expressed genes is shown on the Y-axis.

### Figure S1

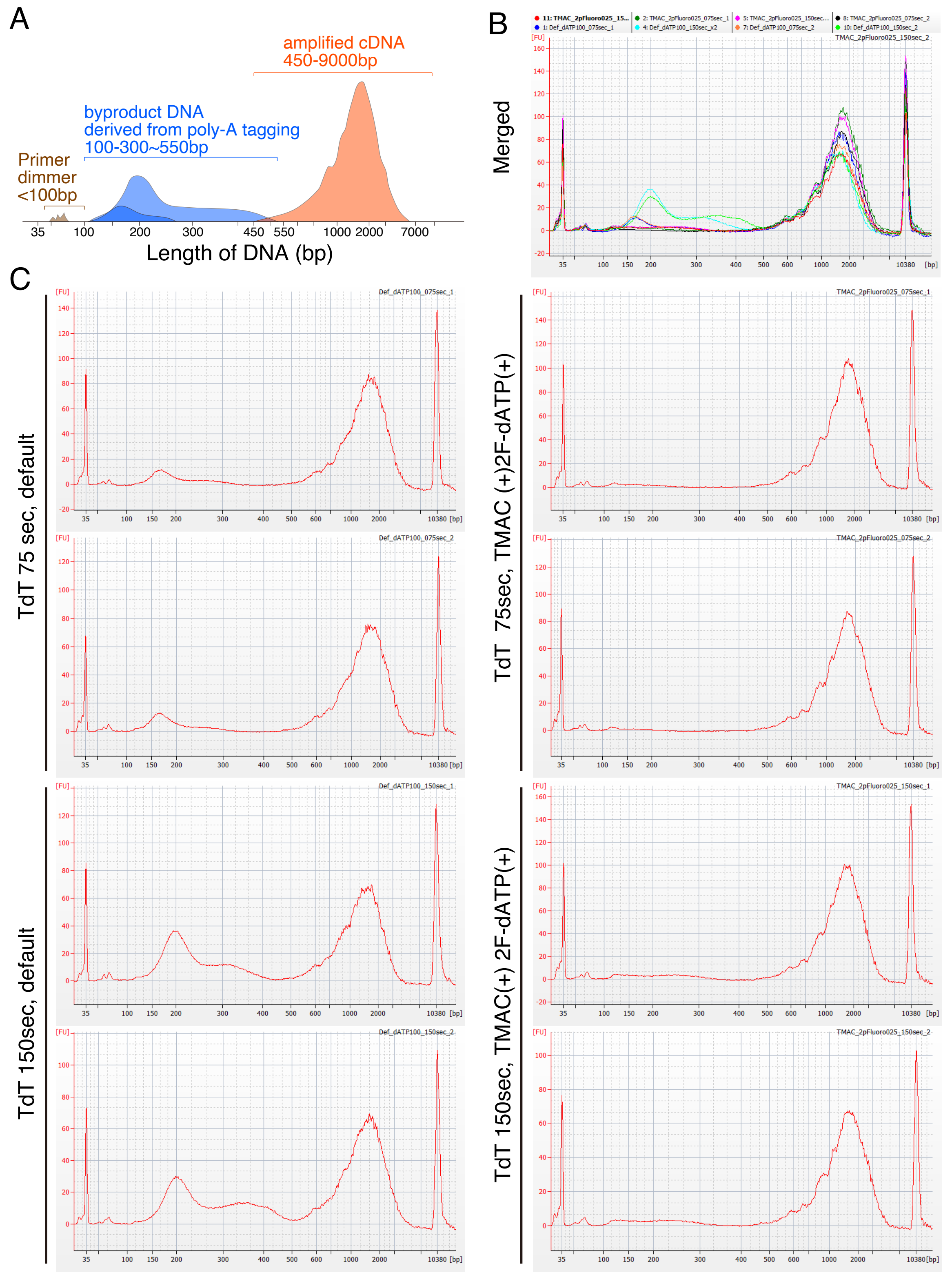

A
amplified cDNA
450-9000bp
byproduct DNA
derived from poly-A tagging
100-300~550bp
Primer
dimmer
<100bp
35
100
200
300
450
550
1000
2000
7000
Length of DNA (bp)
B
Merged
11: TMAC_2pFluoro025_15...
1: Def_dATP100_075sec_1
2: TMAC_2pFluoro025_075sec_1
4: Def_dATP100_150sec_x2
5: TMAC_2pFluoro025_150sec...
7: Def_dATP100_075sec_2
8: TMAC_2pFluoro025_075sec_2
10: Def_dATP100_150sec_2
TMAC_2pFluoro025_150sec_2
[FU]
[bp]
C
TdT 75 sec, default
Def_dATP100_075sec_1
Def_dATP100_075sec_2
TdT 150sec, default
Def_dATP100_150sec_1
Def_dATP100_150sec_2
TdT 75sec, TMAC (+)2F-dATP(+)
TMAC_2pFluoro025_075sec_1
TMAC_2pFluoro025_075sec_2
TdT 150sec, TMAC(+) 2F-dATP(+)
TMAC_2pFluoro025_150sec_1
TMAC_2pFluoro025_150sec_2

### Figure S2

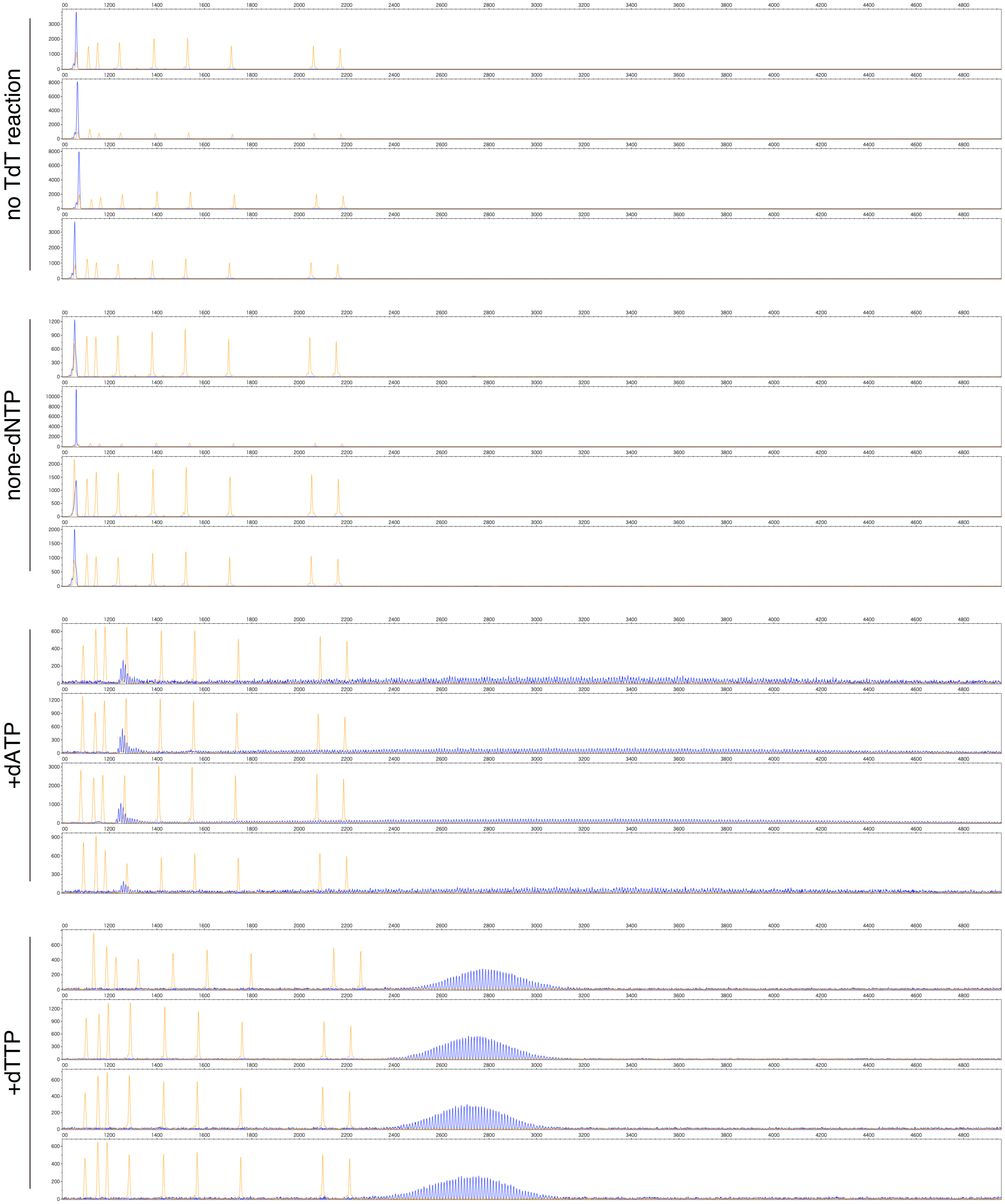

### Figure S3

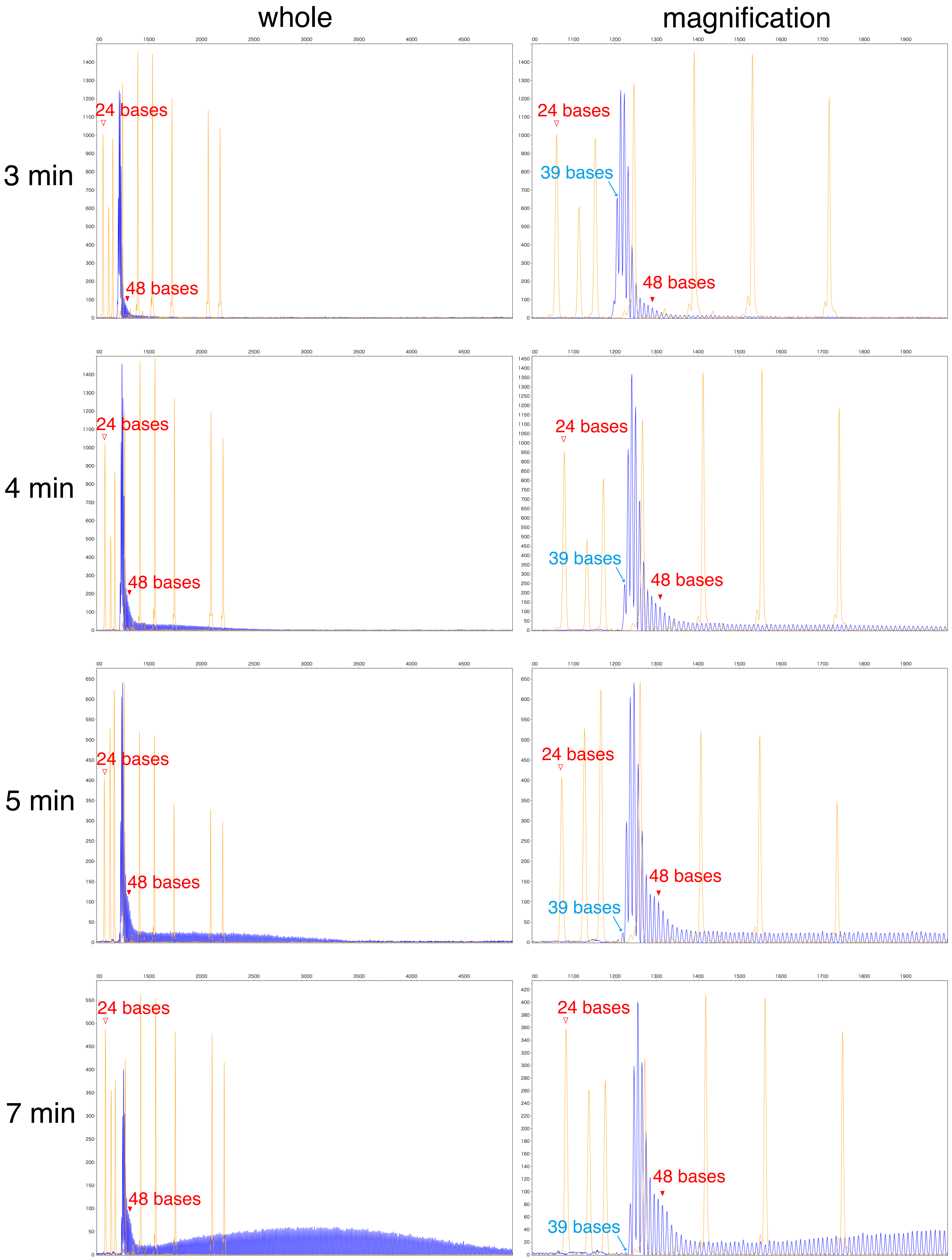

whole
magnification
3 min
4 min
5 min
7 min
24 bases
39 bases
48 bases

### Figure S4

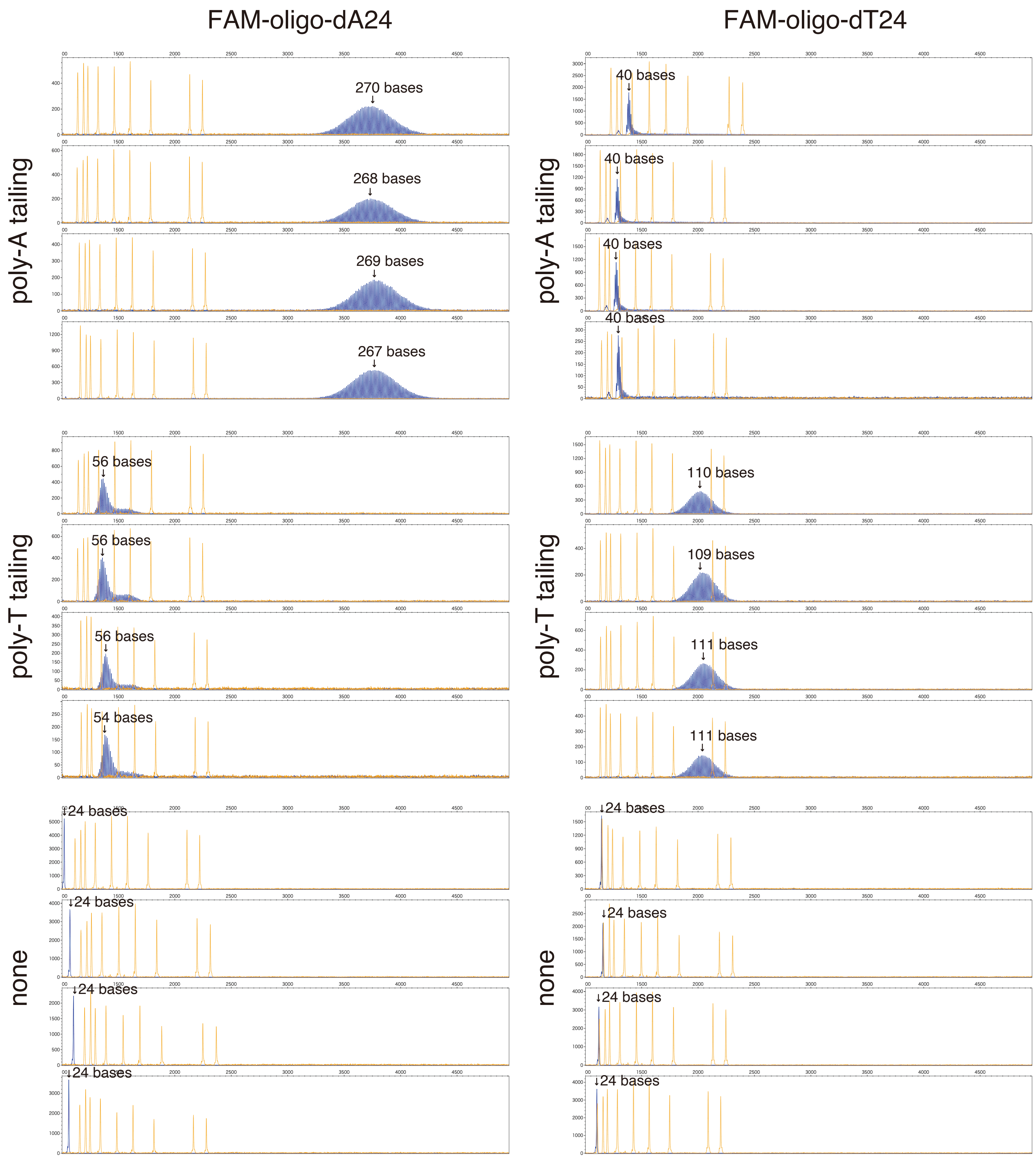
FAM-oligo-dA24
FAM-oligo-dT24
poly-A tailing
poly-T tailing
none
270 bases
268 bases
269 bases
267 bases
40 bases
40 bases
40 bases
40 bases
56 bases
56 bases
56 bases
54 bases
110 bases
109 bases
111 bases
111 bases
24 bases
24 bases
24 bases
24 bases
24 bases
24 bases
24 bases
24 bases

### Figure S5

A

dATP:
2'-Deoxyadenosine
5'-triphosphate

dZTP:
2-Amino-2'
-deoxyadenosine
5'-triphosphate

NH2dATP:
2'-Amino-2'
-deoxyadenosine-
5'-Triphosphate

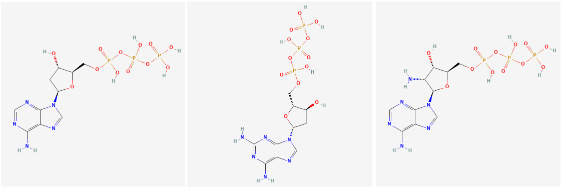

AzdATP:
2'-Azido-2'
-deoxyadenosine-
5'-Triphosphate

OmdATP:
2'-O-Methyladenosine-
5'-Triphosphate

2F-dATP:
2'-Fluoro-2'
-deoxyadenosine
triphosphate

LNA-ATP:
LNA-adenosine-
5'-triphosphate

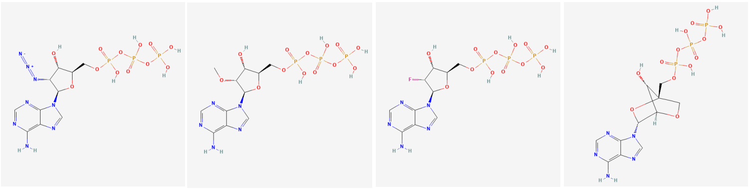

B

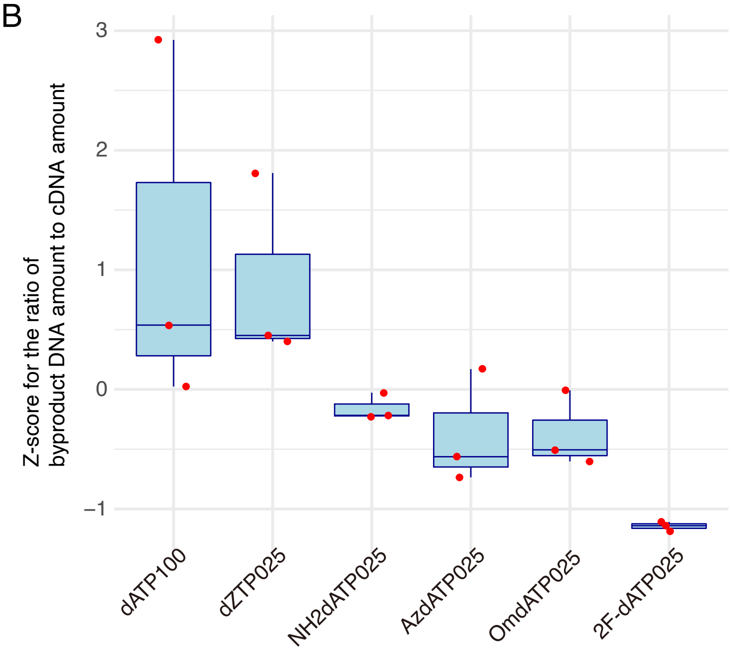


C

dATP (100%)
Default

dZTP (25%)
dATP (75%)

AzdATP (25%)
dATP (75%)

NH2dATP (25%)
dATP (75%)

OmdATP (25%)
dATP (75%)

2F-dATP (25%)
dATP (75%)

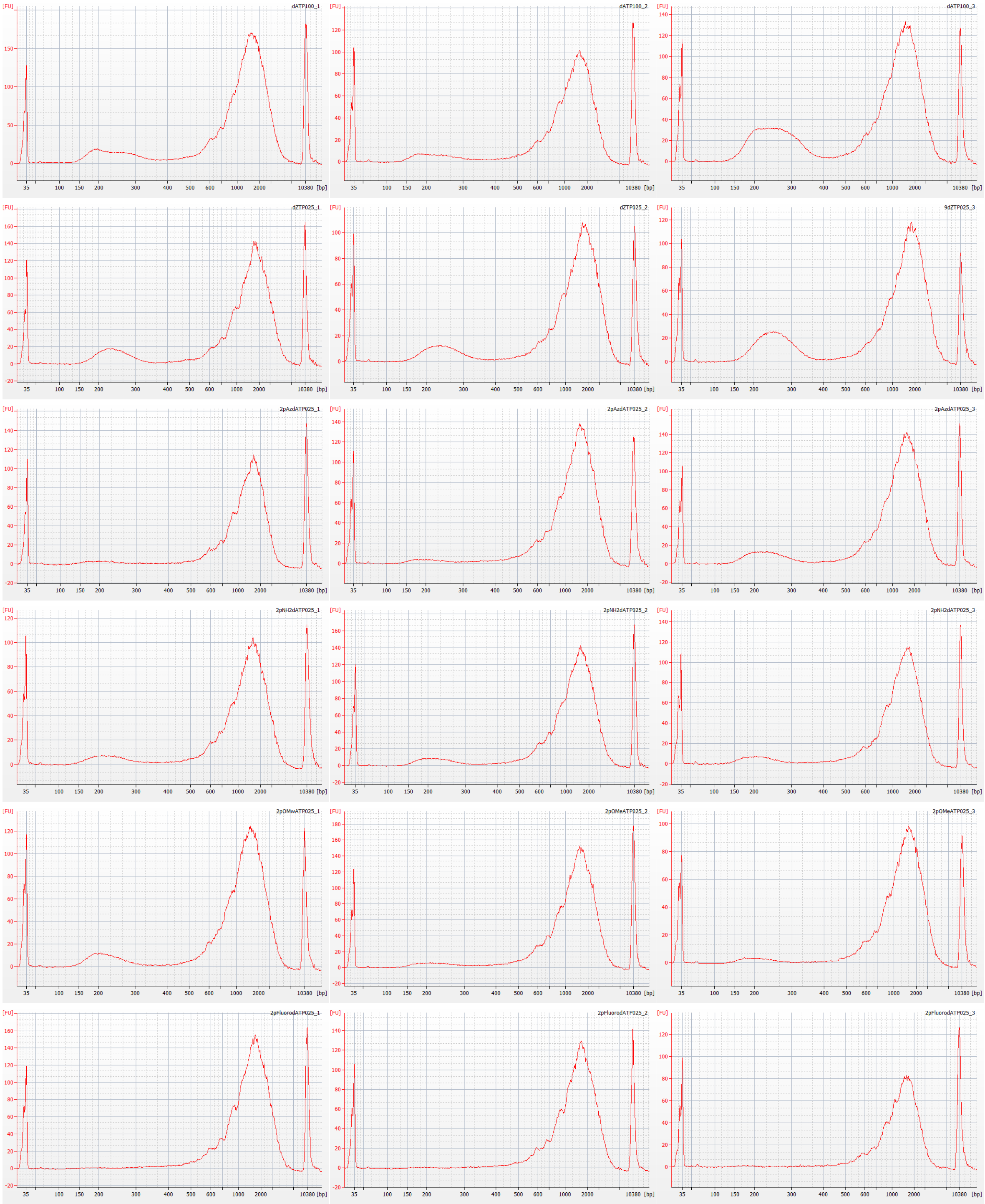

### Figure S6

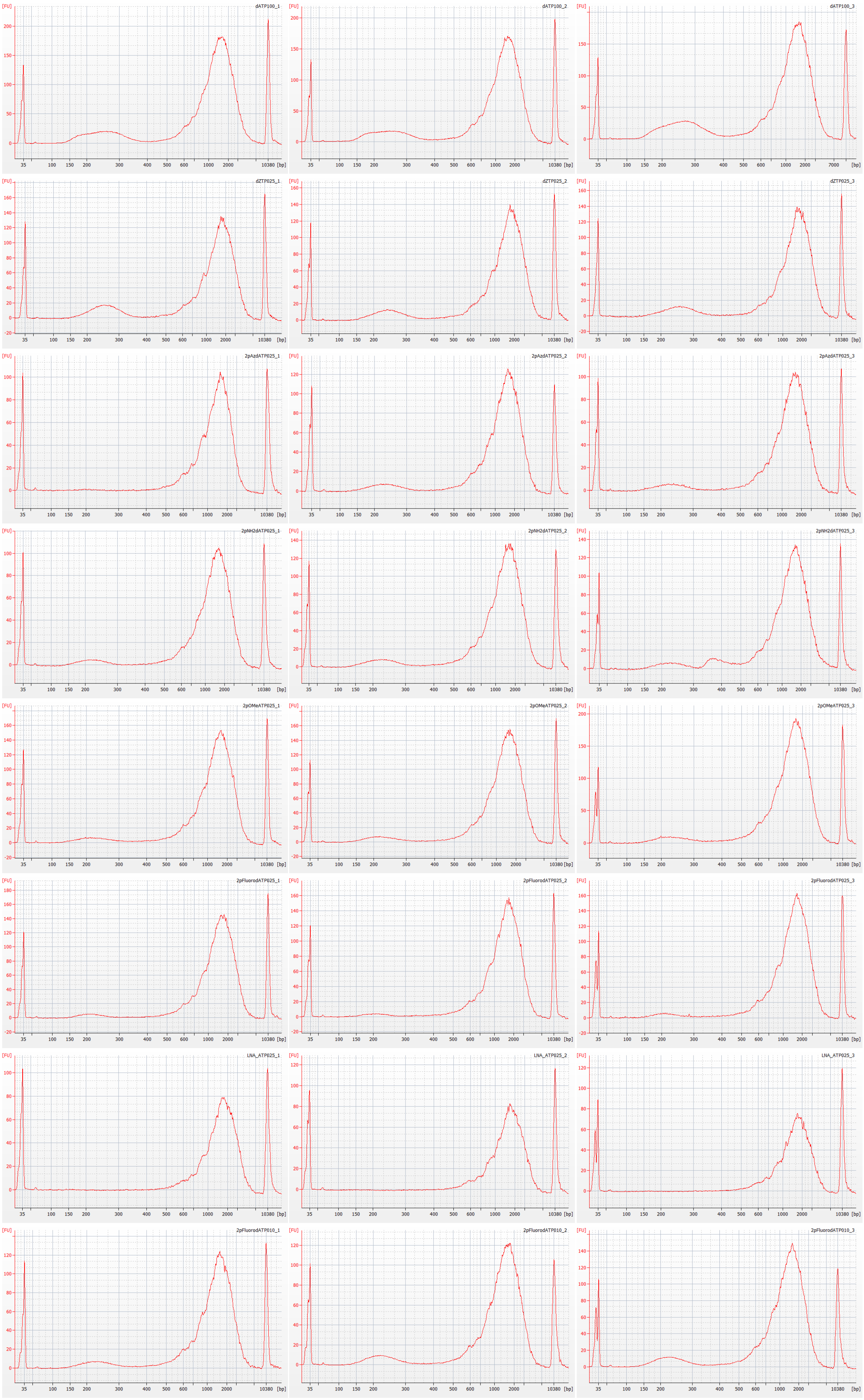

dATP (100%)

dZTP (25%)
dATP (75%)

AzdATP (25%)
dATP (75%)

NH2dATP (25%)
dATP (75%)

OmdATP (25%)
dATP (75%)

2F-dATP (25%)
dATP (75%)

LNA-ATP (25%)
dATP (75%)

2F-dATP (10%)
dATP (90%)

### Figure S7

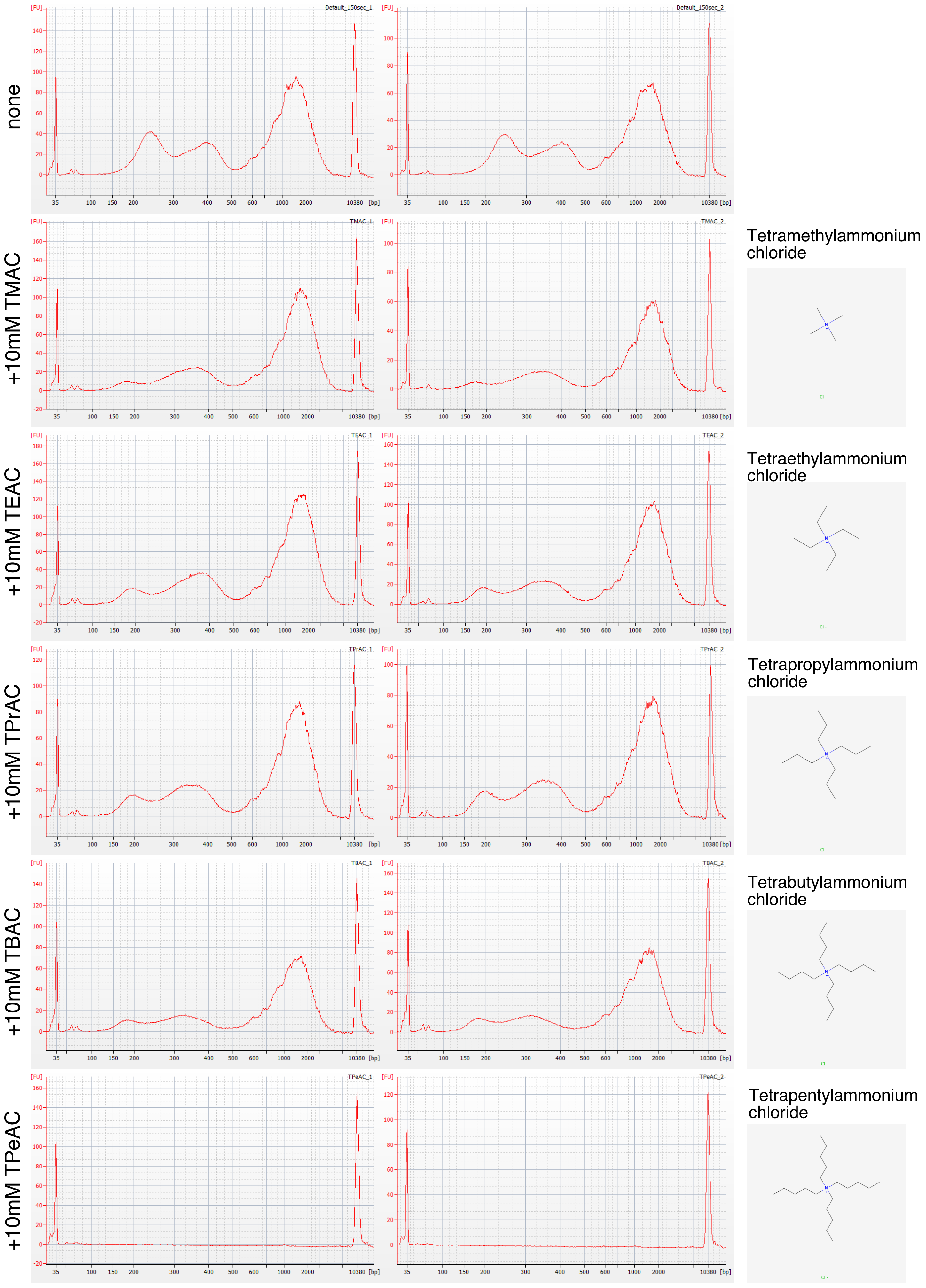
none
Default_150sec_1
Default_150sec_2
+10mM TMAC
TMAC_1
TMAC_2
Tetramethylammonium chloride
+10mM TEAC
TEAC_1
TEAC_2
Tetraethylammonium chloride
+10mM TPrAC
TPrAC_1
TPrAC_2
Tetrapropylammonium chloride
+10mM TBAC
TBAC_1
TBAC_2
Tetrabutylammonium chloride
+10mM TPeAC
TPeAC_1
TPeAC_2
Tetrapentylammonium chloride
[FU]
[bp]
35
100
150
200
300
400
500
600
1000
2000
10380

### Figure S8

A

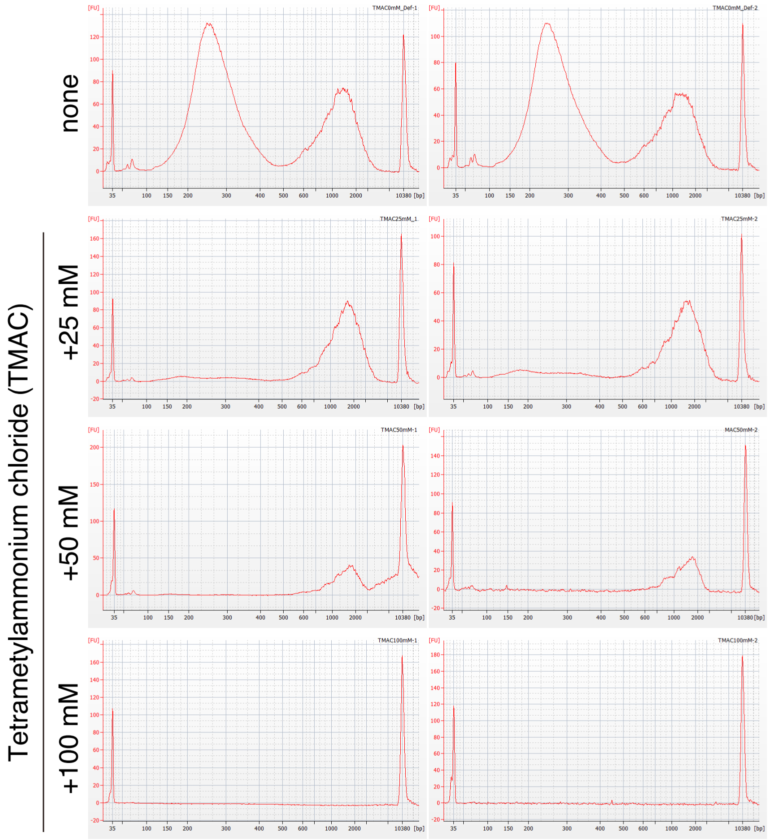


B

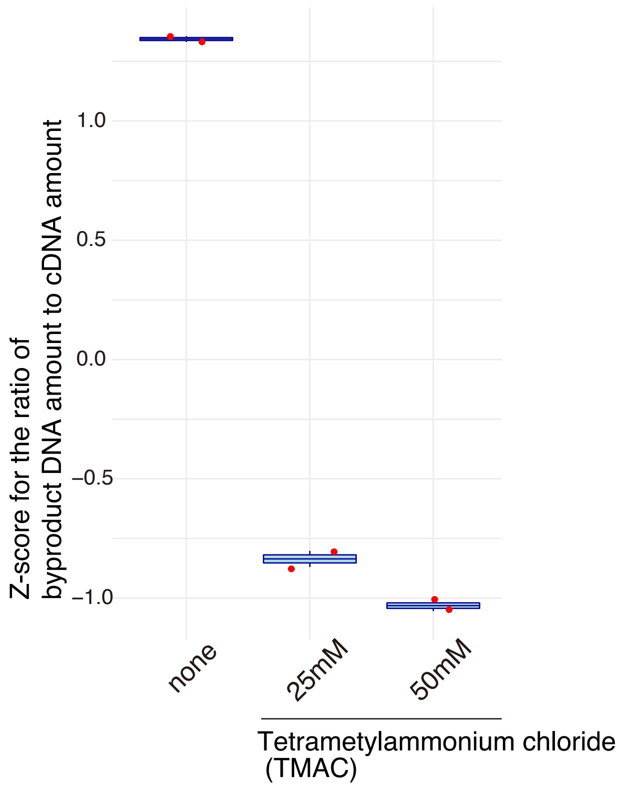


C

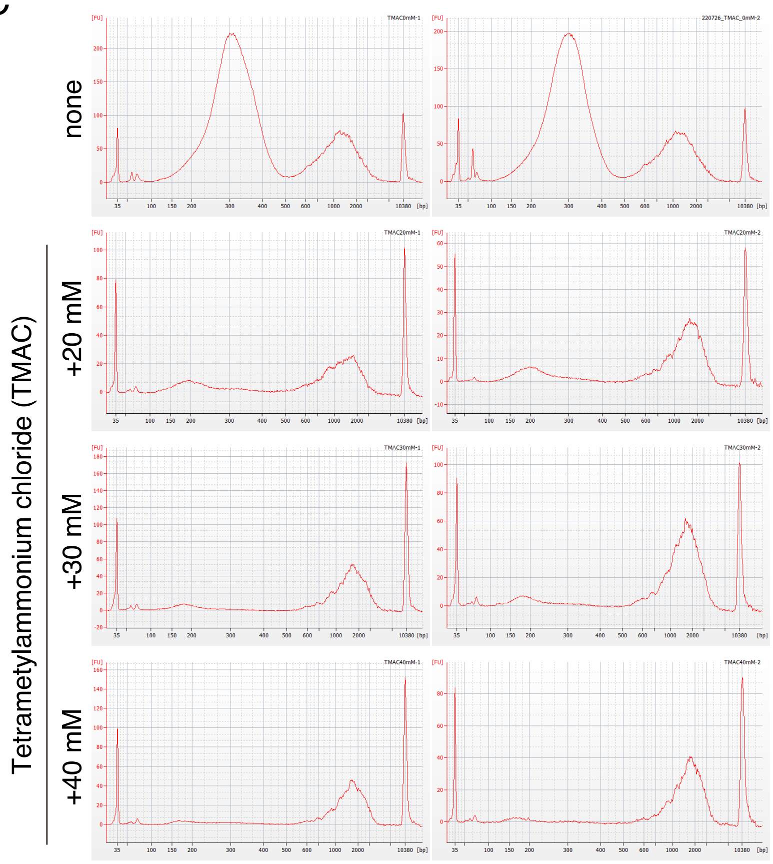


D

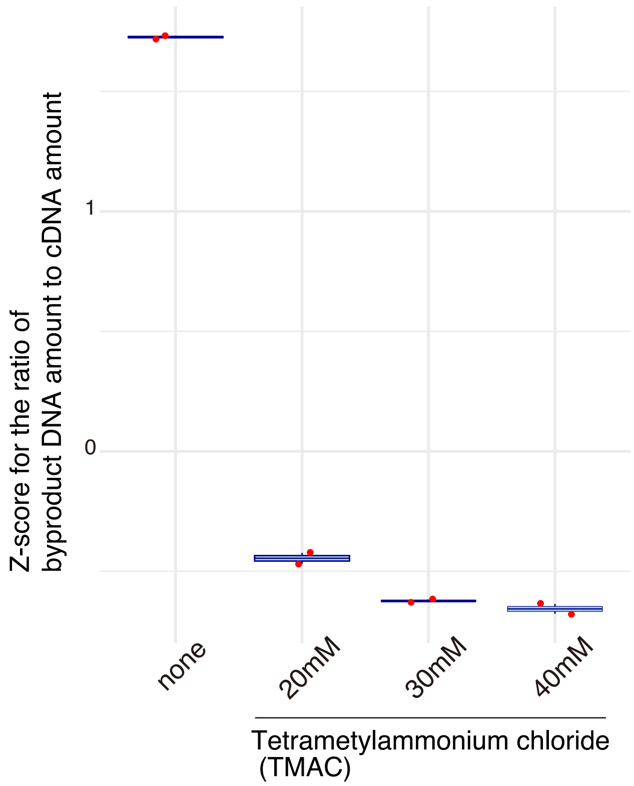

### Figure S9

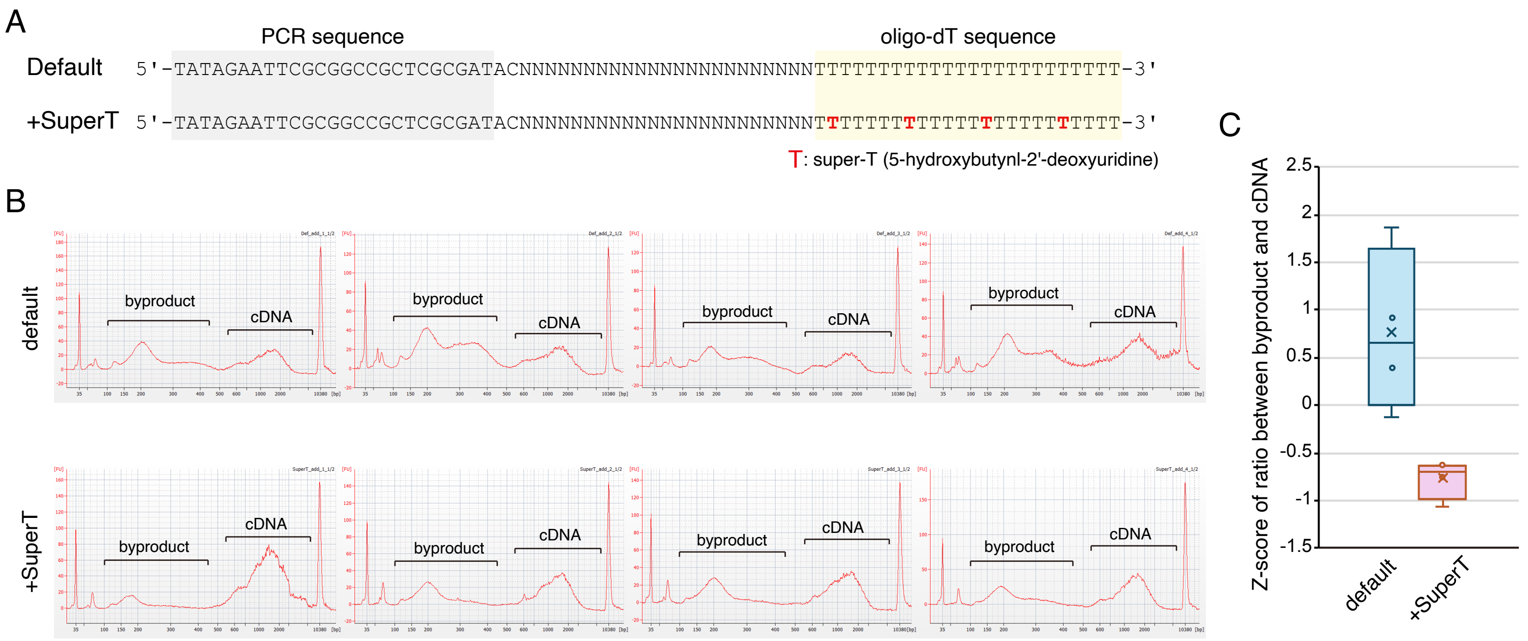
A
PCR sequence
oligo-dT sequence
Default 5'-TATAGAATTCGCGGCCGCTCGCGATACNNNNNNNNNNNNNNNNNNNNNNNTTTTTTTTTTTTTTTTTTTTTTTT-3'
+SuperT 5'-TATAGAATTCGCGGCCGCTCGCGATACNNNNNNNNNNNNNNNNNNNNNNNTTTTTTTTTTTTTTTTTTTTTTTT-3'
T: super-T (5-hydroxybutynl-2'-deoxyuridine)
B
default
+SuperT
byproduct
cDNA
C
Z-score of ratio between byproduct and cDNA
2.5
2
1.5
1
0.5
0
-0.5
-1
-1.5
default
+SuperT

### Figure S10

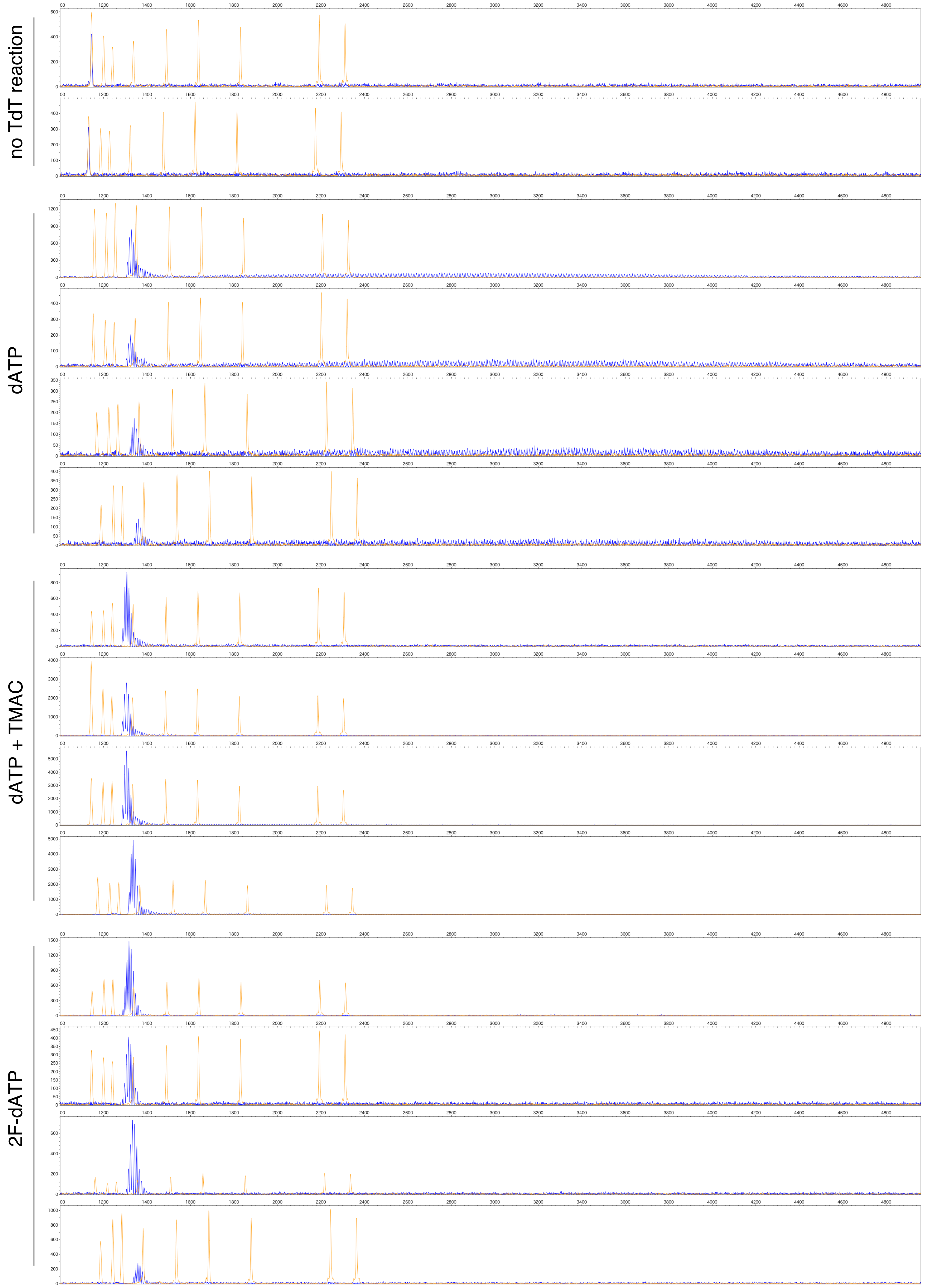

### Figure S11

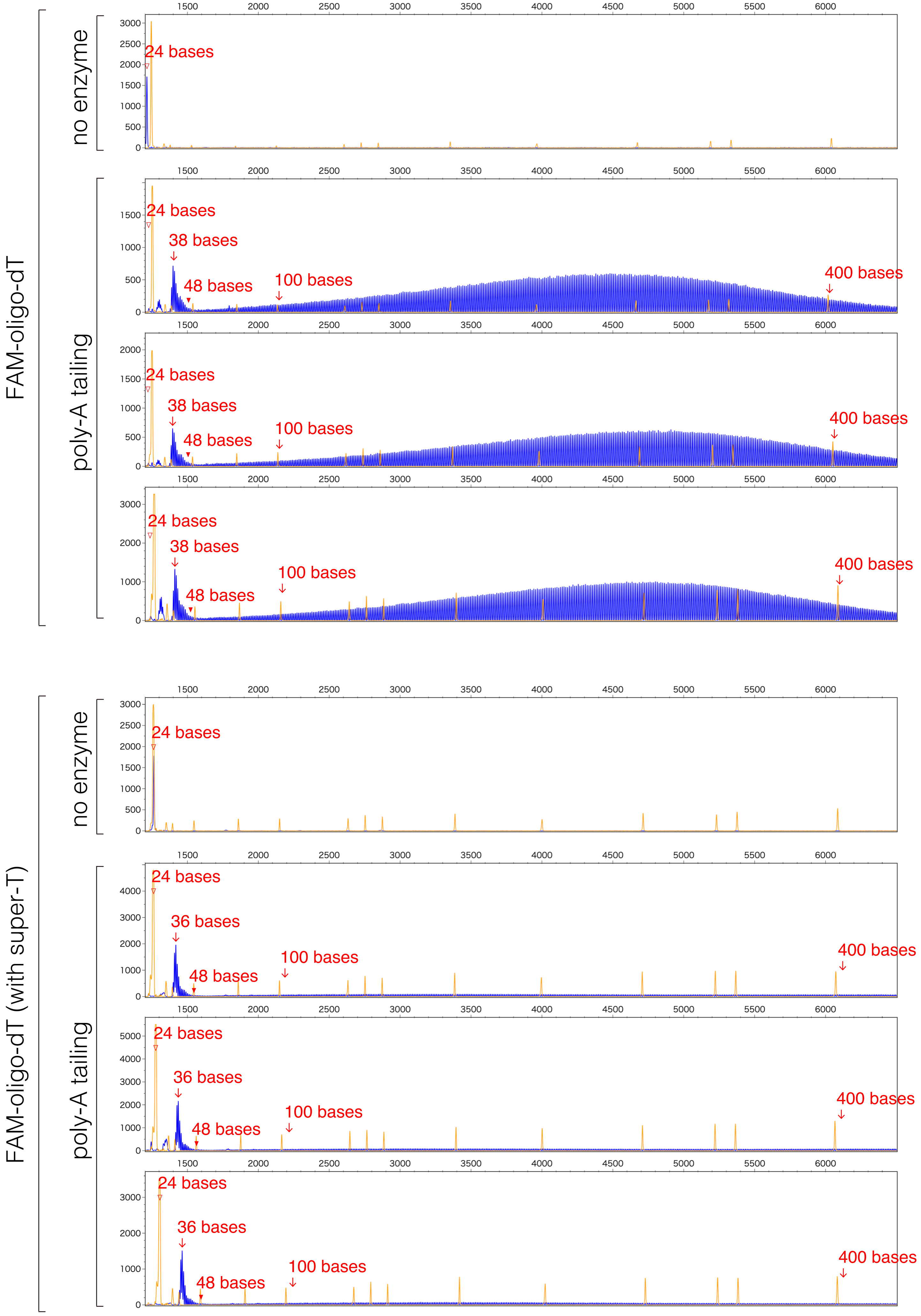

FAM-oligo-dT
no enzyme
poly-A tailing
24 bases
38 bases
48 bases
100 bases
400 bases
FAM-oligo-dT (with super-T)
36 bases

Figure S12

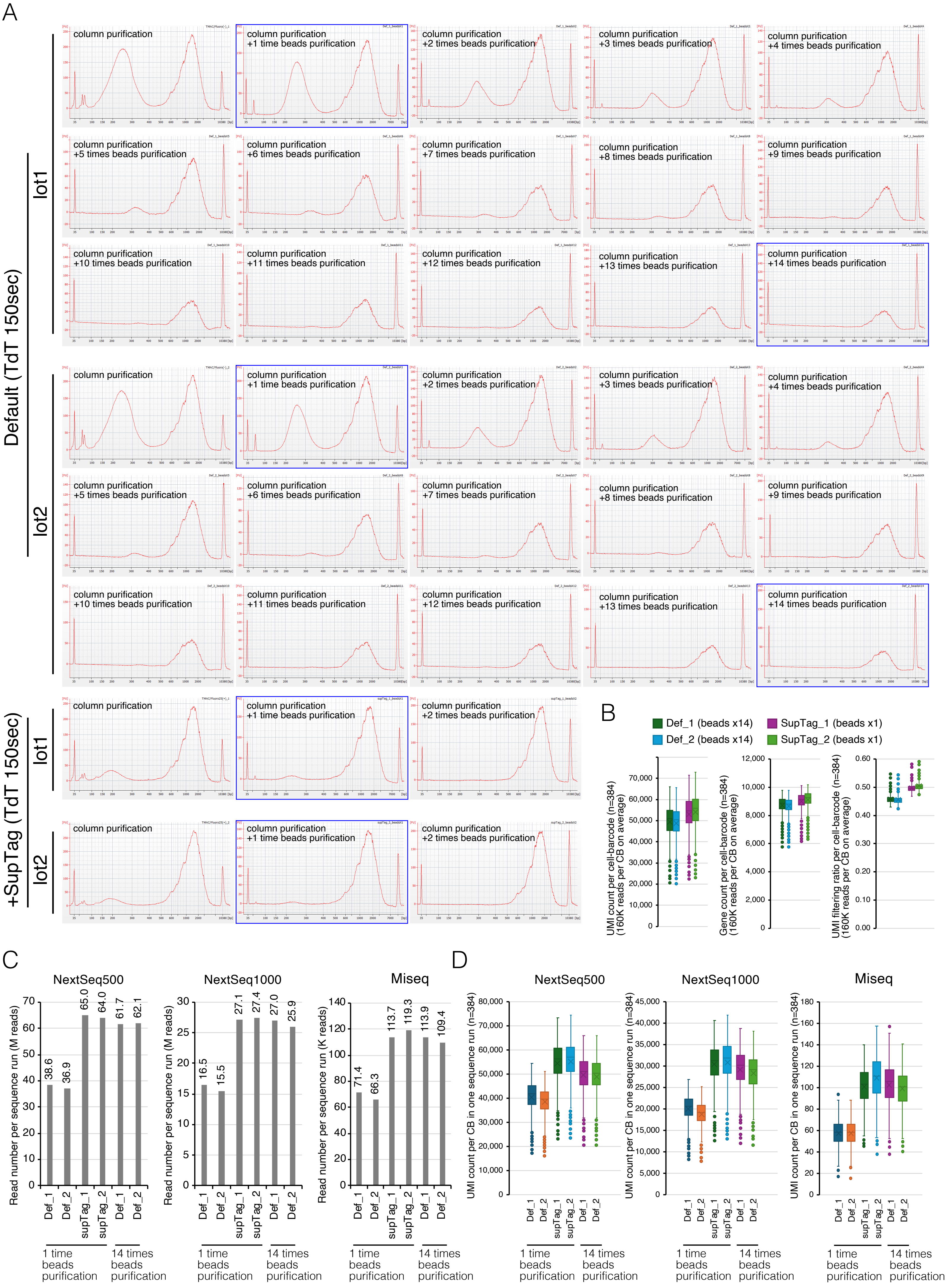
A
Default (TdT 150sec)
lot1
lot2
+SupTag (TdT 150sec)
column purification
column purification +1 time beads purification
column purification +2 times beads purification
column purification +3 times beads purification
column purification +4 times beads purification
column purification +5 times beads purification
column purification +6 times beads purification
column purification +7 times beads purification
column purification +8 times beads purification
column purification +9 times beads purification
column purification +10 times beads purification
column purification +11 times beads purification
column purification +12 times beads purification
column purification +13 times beads purification
column purification +14 times beads purification
B
Def_1 (beads x14)
Def_2 (beads x14)
SupTag_1 (beads x1)
SupTag_2 (beads x1)
UMI count per cell-barcode (n=384) (160K reads per CB on average)
Gene count per cell-barcode (n=384) (160K reads per CB on average)
UMI filtering ratio per cell-barcode (n=384) (160K reads per CB on average)
C
NextSeq500
NextSeq1000
Miseq
Read number per sequence run (M reads)
Read number per sequence run (K reads)
38.6
36.9
65.0
64.0
61.7
62.1
16.5
15.5
27.1
27.4
27.0
25.9
71.4
66.3
113.7
119.3
113.9
109.4
Def_1
Def_2
supTag_1
supTag_2
1 time beads purification
14 times beads purification
D
NextSeq500
NextSeq1000
Miseq
UMI count per CB in one sequence run (n=384)

### Figure S13

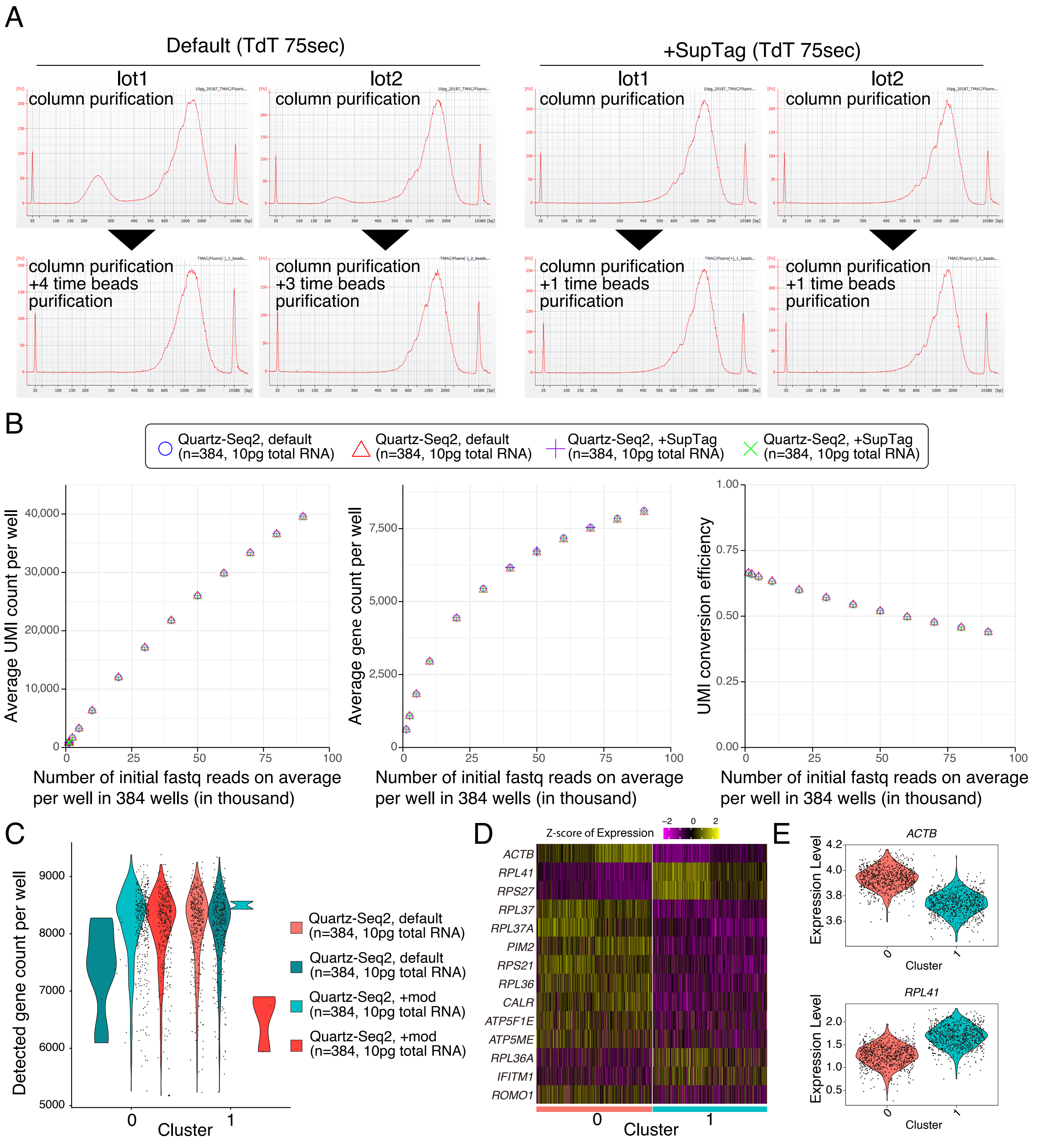

A
Default (TdT 75sec)
+SupTag (TdT 75sec)
lot1
lot2
column purification
column purification +4 time beads purification
column purification +3 time beads purification
column purification +1 time beads purification
B
Quartz-Seq2, default (n=384, 10pg total RNA)
Quartz-Seq2, +SupTag (n=384, 10pg total RNA)
Average UMI count per well
Average gene count per well
UMI conversion efficiency
Number of initial fastq reads on average per well in 384 wells (in thousand)
C
Detected gene count per well
Cluster
Quartz-Seq2, default (n=384, 10pg total RNA)
Quartz-Seq2, +mod (n=384, 10pg total RNA)
D
Z-score of Expression
ACTB
RPL41
RPS27
RPL37
RPL37A
PIM2
RPS21
RPL36
CALR
ATP5F1E
ATP5ME
RPL36A
IFITM1
ROMO1
Cluster
E
ACTB
RPL41
Expression Level
Cluster
